# Sequence, Structural and Functional Diversity of the Ubiquitous DNA/RNA-Binding Alba Domain

**DOI:** 10.1101/2024.04.16.589850

**Authors:** Jagadeesh Jaiganesh, Shruthi Sridhar Vembar

## Abstract

The DNA/RNA-binding Alba domain is prevalent across all kingdoms of life. Discovered in archaea more than 40 years ago, this protein domain appears to have evolved from RNA- to DNA- binding, with a concomitant expansion in the range of cellular processes that it regulates. Indeed, Alba domain-containing proteins continue to exhibit functional ingenuity, with their most recent association being R-loop regulation in plants: they accomplish this by binding to DNA:RNA hybrid structures as heterodimers. To further explore Alba diversity and evolutionary relatedness, which could in turn lead to new functional revelations, we employed iterative searches in PSI-BLAST to identify 15161 true Alba domain-containing proteins from the NCBI non-redundant protein database. Next, by building sequence similarity networks (SSNs), we identified 13 distinct clusters representing various Alba subgroups. These included the classic eukaryotic Rpp20/Pop7 and Rpp25/Pop6 proteins as well as novel fungal Alba proteins and *Plasmodium*-specific Albas. Conservation analysis indicated that, despite overall low primary sequence similarity between the SSN clusters, the homo- or hetero-dimer interface is highly conserved. Furthermore, the presence of the classic Alba fold was confirmed in representative sequences from each cluster by comparison of their Hidden Markov Model (HMM) profiles and *ab initio* three-dimensional structures. Notably, Alba domains from lower, unicellular eukaryotes and fungal Albas exhibit structural deviations towards their C-terminal end. Finally, phylogenetic analysis, while supporting SSN clustering, revealed the evolutionary branchpoint at which the eukaryotic Rpp20- and Rpp25-like clades emerged from archaeal Albas, and the subsequent taxonomic lineage-based divergence within each clade. Taken together, this comprehensive analysis enhances our understanding of the evolutionary history of Alba domain-containing proteins across diverse organisms, their sequence and structural conservation, and functional implications for genome and RNA biology.

## Introduction

Genome organization and gene regulation are intertwined and indispensable processes in all domains of life. In eukaryotes, these processes are largely controlled by DNA-binding proteins, histones and their regulators, while in prokaryotes, several nucleoid-associated proteins (NAPs) take up this role. NAPs can bend, bridge, wrap, and polymerize along DNA, which ultimately allows them to exert a pervasive effect on gene expression. In addition, NAPs involved in sequence-independent chromatin organization are hypothesized to take on the role of transcription factors at certain genetic loci (reviewed in Visweswariah & Busby, 2015). One such NAP found in all archaeal species is called Alba (Acetylation Lowers Binding Affinity): as the name suggests, Alba can exist in acetylated and non-acetylated states, with the former binding to DNA with decreased affinity (Bell et al., 2002). Since their initial discovery in archaea in the 1980s (Lurz et al., 1986), Alba-domain containing proteins have been described in a variety of lower and higher eukaryotic organisms (Paysan-Lafosse et al., 2023), with most organisms expressing a minimum of two Alba domain-containing proteins. Moreover, with the advent of whole genome sequencing and a concomitant increase in the number of reported protein sequences, additional domains similar to Alba have been reported, leading to the classification of an “Alba-like domain superfamily” in the InterPro database (IPR036882), which includes Rpp20/Pop7 and Rpp25/Pop6 protein components of the RNAse P/MRP complex, Sporulation stage V, protein S (SpoVS) of bacteria, and other Alba domain-containing proteins.

Phylogenetic analyses suggested that Alba originated as an RNA-binding domain in archaea (Aravind et al., 2003), and with time, evolved to bind to DNA and regulate chromatin biology (Dijk & Reinhardt, 1986). The crystal structure of an archaeal Alba homodimer revealed a domain architecture of two α-helices and four β-strands arranged in β1-α1-β2-α2-β3-β4 configuration, which is similar to the ancient IF3-C fold found at the C-terminal end of RNA-binding translation initiation factors, as well as to the fold of DNAse I (Aravind et al., 2003; Wardleworth et al., 2002). Within archaea, Alba proteins show functional diversity: in mesophilic archaea, they are thought to be transcription factors (Liu et al., 2009), while in euryarchaeota, they are shown to bind to RNA and potentially serve as RNA-stabilising or -folding proteins or as translational regulators (L. Guo et al., 2014; R. Guo et al., 2003; N. Zhang et al., 2020). Moreover, given that many archaeal species encode divergent Alba domain-containing proteins Alba1 and Alba2, with sequence identities as little as 30%, this protein family can form both homo- and hetero- dimers, each packaging DNA and/or RNA into higher order structures in a distinct manner (Jelinska et al., 2005, 2010). Nonetheless, Alba-nucleic acid binding is enabled through contacts between cationic amino acid residues in the Alba multimer and sugar phosphates in the DNA or RNA backbone (L. Guo et al., 2014; Tanaka et al., 2012).

Early on, Alba domain proteins garnered interest as putative therapeutic targets in unicellular eukaryotic pathogens. For instance, the malaria-causing Apicomplexan parasite *Plasmodium falciparum* encodes for six Alba-domain containing proteins (Chêne et al., 2012; Goyal et al., 2012; Reddy et al., 2015). Of these, PfAlba1, PfAlba2, PfAlba3 and PfAlba4 bind to both DNA and RNA (Chêne et al., 2012), and an in-depth characterization of PfAlba1 showed its role in fine-tuning the timing of translation of virulence factor-encoding mRNAs during asexual blood stage development (Vembar et al., 2015). PfAlba3 has independently been implicated in regulating the expression of sub-telomeric *var* genes, which encode for virulence-associated surface antigens required for *P. falciparum* immune evasion (Goyal et al., 2012), and as an endonuclease *in vitro* (Banerjee et al., 2023). In contrast, very little is known about PfAlba5 and PfAlba6, which were discovered more recently using an *in silic*o approach and exhibit less than 15% sequence similarity to the remaining PfAlbas (Reddy et al., 2015). In addition to *Plasmodium sp.,* Alba domain-containing proteins from the Discoban parasites *Trypanosoma brucei* and *Leishmania infantum* have been implicated as master regulators of proteome remodelling during stage transitions (Bevkal et al., 2021; da Costa et al., 2017; Ferreira et al., 2020) while Alba proteins from *Toxoplasma gondii* are involved in translational control of gene expression (Gissot et al., 2013).

In other uni- and multi-cellular eukaryotes, Alba domain-containing proteins have been divided into two major families, namely the Ribonuclease P protein subunit p20 family (Rpp20; also called Pop7 in budding yeast *Saccharomyces cerevisiae*) and RNase P protein subunit p25 family (Rpp25; also called Pop6). Rpp20/Pop7 and Rpp25/Pop6 were initially shown to interact as a heterodimer with the P3 domain of the catalytic RNA of the RNase P/MRP holoenzyme, which is an essential regulator of tRNA maturation in all domains of life (Chan et al., 2018; Yin et al., 2021). Besides, these subunits can form homodimers *in vitro*, although the functional relevance of homodimerization is not known (Hands-Taylor et al., 2010). The individual subunits also function independently; for example, human Rpp20 has ATPase activity (Li & Altman, 2001) and complexes with the heat shock protein Hsp27 which is known to enhance the activity of RNase P *in vitro* (Jiang & Altman, 2001). More recently, it was shown that the binding of *S. cerevisiae* Pop6-Pop7 to the telomerase RNA as a heterodimer stabilizes the binding of other telomerase components, thus making them essential constituents of this ribonucleoprotein complex (Lemieux et al., 2016). In plants, multiple Alba-domain containing proteins are present but their functions are less explored. Best studied in *Arabidopsis thaliana,* AtAlba1 and AtAlba2 are genic R-loop readers which maintain genome stability while AtAlba4 and AtAlba6 are involved in RNA metabolism, male reproductive development and heat stress response (Náprstková et al., 2021; Yuan et al., 2019). Stress-induced expression of Alba proteins has also been observed in *Oryza sativa* (Verma et al., 2014). Overall, the functional diversity exhibited by Alba domain-containing proteins across different kingdoms of life hints at the evolutionary capability and the ‘innovability’ of this protein domain.

In this study, to understand the mechanisms by which the Alba domain evolved to gain divergent functions, we performed a comprehensive sequence–structure–function relationship analysis of more than 15000 Alba domain-containing proteins from archaea, bacteria and eukaryotes. By combining sequence similarity networks (SSNs), multiple sequence alignments (MSAs), Hidden Markov Model (HMM) profiles, sequence conservation scores, *de novo* structural comparisons, and phylogenetic analysis, we identified thirteen distinct clusters of Alba domain-containing proteins and determined their evolutionary relatedness to each other. Intriguingly, many of these clusters are specific to certain unicellular eukaryotic lineages such as *Plasmodium, Theileria, Saccharomycetales* and other fungal groups. In addition, our analysis reclassified select Alba family members from pathogenic eukaryotes such as *Plasmodium sp.* into clades divergent from their host organism, reiterating their therapeutic potential.

## Materials and Methods

### Gathering Alba-domain containing proteins

A list of Alba domain-containing proteins from various domains of life was compiled from literature searches, and the sequences of their Alba domains extracted in FASTA format; the average length of the domain was found to be 90 to 120 amino acids in archaea, and 100 to 160 amino acids in eukaryotes. Homology modeling was used to confirm that the extracted sequences corresponded solely to the Alba fold, which were then used as ‘seed’ sequences for preliminary searches in PSI-BLAST (Altschul et al., 1997) against the NCBI non-redundant (NR) protein database (Sayers et al., 2022) with an E-value cut-off of 10 and a total of five iterations. The resulting list of ‘hits’ was filtered at 90% maximum pairwise identity using MMseqs2 (Steinegger et al., 2021) and clustered using CLANS (Frickey & Lupas, 2004) with default parameters. Clusters were defined based on a p-value cutoff of 1e-4. For clusters that did not contain any of the initial seed sequences, a consensus sequence was created and the ‘hit’ with highest sequence identity to the consensus was chosen as the best representative. These representative sequences were manually investigated to confirm the presence of the Alba domain using HHpred searches (Söding et al., 2005) and RoseTTAFold structure predictions (Baek et al., 2021). Clusters that passed these investigations were retained and their representative sequences used for a second round of PSI-BLAST; all other outliers in the CLANS cluster map were removed. This process was iterated until no new clusters were discovered. The resulting final list of seed sequences was used to query the September 2021 release of the NCBI NR protein database by performing PSI-BLAST with an E-value cut-off of 10 and a total of five iterations. The hits were clustered in CLANS after filtering for a maximum pairwise identity of 90% using MMSeqs2 and clusters defined based on a p-value cutoff of 1e-4. The Krona visualisation tool (Ondov et al., 2011) was used to examine the extent of distribution of Alba domain-containing proteins across all taxonomic groups and in each cluster.

### Network building and visualisation

The Enzyme Function Initiative’s (EFI’s) web resources (Zallot et al., 2019) were used to generate the Alba Sequence Similarity Network (SSN), which was built by performing an all-against-all BLAST to calculate the pairwise similarities of all input sequences. A FASTA file containing 8390 sequences was used as input, and an E-value of 4 was used to calculate the SSN edge alignment score similarities, as described in the web resources (https://efi.igb.illinois.edu/efi-est/). Because visualising the entire network was computationally expensive, a representative network filtered to a maximum pairwise identity of 55 percent using CD-HIT (Huang et al., 2010) and containing 5,967 nodes and 2,033,164 edges was generated. For network visualisation, the Prefuse force-directed layout in Cytoscape v3.9.1 (Shannon et al., 2003) was used, with edge E-value threshold set to 1E-4. Clusters were defined based on the CLANS map shown in **Supplementary Figure S1**.

### Identifying distant homologs using HHpred and pairwise HMM comparison

Using the final list of 31 Alba seed sequences as input, HHpred searches were performed in January 2022 against the PDB mmCIF70, SCOPe70, Pfam-A v35, and ECOD F70 databases, as well as proteomes of some model organisms whose profile HMMs were available in the MPI Bioinformatics toolkit database (Gabler et al., 2020). This ensured that the searches had converged and that all members of the Alba superfamily as well as distant homologs had been assimilated. Next, for seed sequences and the newly identified distant homologs, pairwise sequence similarity was calculated using the HHalign module in HH-suite3 (Steinegger et al., 2019). To create profile HMM for each sequence, three iterations of HHblits (Steinegger et al., 2019) was run against the Uniclust30 database (Mirdita et al., 2017) with an E-value inclusion threshold of 1E-3. The profile HMMs were then pairwise aligned with HHalign to obtain the HHpred probability percentage.

### Protein Structure Prediction and comparison

RoseTTAFold, an accurate protein structure prediction method based on deep learning (Baek et al., 2021), was used to predict the structures of the 31 seed sequences and three distant homologs *Thermus thermophilus* SpoV (PDB ID:2EK0; Rehse & Yokoyama, 2007), *Escherichia coli* YbhY (PDB ID:1LN4; Ostheimer et al., 2002) and *E. coli* IF3-C (PDB ID:2IFE; De Cock et al., 1998). Although crystal structures for some of the seed sequences were available, all structures were predicted using RoseTTAFold to ensure uniformity. TM-scores were calculated using US-align (C. Zhang et al., 2022) for pairwise comparisons of all predicted structures, and the accuracy of the predicted structures was also compared to crystal structures wherever available. All models were visualised in PyMol 2.5.2 (Schrödinger, LLC, 2015), retrieved from http://www.pymol.org/pymol. For all predicted structures, the electrostatic potential surface charges were calculated using the APBS Electrostatic Potential plugin (Baker et al., 2001) in PyMol.

### Sequence Conservation and Phylogenetic analysis

All sequence alignments were performed using the MAFFT v.7.490 program (Katoh et al., 2019) with the L-INS-i algorithm. Conservation of amino acid residues was quantified using the Scorecons server (Valdar, 2002) against the sequences whose crystal structures were available (Chan et al., 2018; Liu et al., 2009; Kumarevel et al., 2008; Perederina et al., 2007; Zhao et al., 2003). For the phylogenetic reconstruction, sequences from each cluster in the CLANS cluster map, except for clusters with fewer than 30 sequences, were extracted and filtered using MMseqs2 at a maximum pairwise identity of 70% to obtain a list of 4666 sequences. The alignment was then trimmed using Goalign ‘clean sites’ option (Fŕ et al., 2021). A maximum likelihood tree was inferred using IQTree v2.1.2 (Minh et al., 2020) with 3000 bootstrap iterations and automatic substitution model selection (Kalyaanamoorthy et al., 2017). For in-depth inferences, -nstop was set to 300 and -pers to 0.3. iTOL (Letunic & Bork, 2021) was used to visualise and annotate the tree.

## Results

### Alba proteins are found in all domains of life

To obtain a comprehensive picture of the diversity of Alba domain-containing proteins, we chose to implement iterative sequence similarity searches in PSI-BLAST (Altschul et al., 1997) against the NCBI non-redundant (nr) protein sequence database (Sayers et al., 2022), similar to a previous report (Alva & Lupas, 2019). To begin with, ‘seed’ sequences corresponding to well-characterised Alba domain-containing proteins described in the literature from archaea, unicellular eukaryotes, plants, fungi, and multicellular eukaryotes, and bacterial protein sequences annotated as ‘Alba-domain containing’ in the NCBI nr protein database, were used as queries in PSI-BLAST. When the resulting hits were clustered and visualized using CLANS (Frickey & Lupas, 2004), two new clusters which did not contain any of the initial seed sequences were identified. After confirming that these clusters were truly Alba-like, representative sequences from these were used as queries in PSI-BLAST against the nr database, and clustered and visualised in CLANS, resulting in the identification of three new clusters; this process was continued until no new clusters were formed. Finally, 31 Alba domain sequences (**Table 1**) were used as ‘seeds’ in PSI-BLAST against the NCBI nr protein database to identify Alba proteins from all domains of life.

**Table 1.**
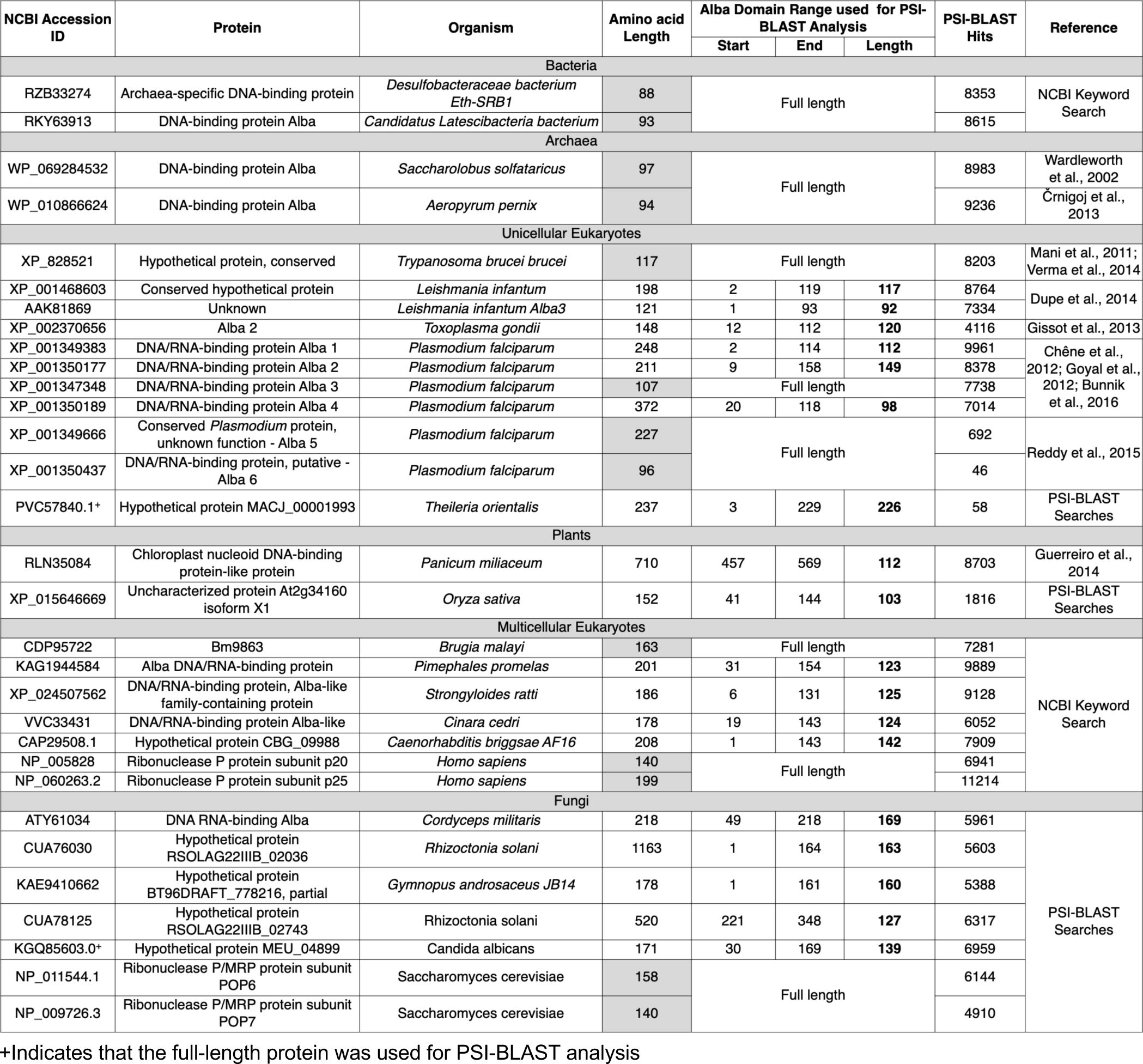
List of Alba seed sequences used for PSI-BLAST analyses. The table lists 31 Alba domain-containing proteins shortlisted for PSI-BLAST through literature survey, keyword searches in NCBI and preliminary cluster analysis using CLANS.

The resulting 16547 unique hits (**Figure 1A**) were filtered to retain true Alba-domain containing proteins. First, 42 isoforms of the 4.2 MDa protein Titin were removed from the list since, upon closer scrutiny, they did not contain the signature Alba fold. Next, MMseqs2 (Steinegger et al., 2021) was used to reduce sets of closely related sequences at a maximum pairwise identity of 90% to a single representative sequence, resulting in 9197 entries. Lastly, with the help of CLANS cluster maps, outlier sequences which either remained as singletons, or clustered without forming edges with seed clusters at a pairwise BLAST p-value threshold of 1e-4, or lacked an Alba domain, were removed (**Figure 1A**). The final CLANS cluster map (**Supplementary Figure S1**) contained 8390 sequences which were mapped back to the original 90% MMSeqs2-reduced list, to obtain a final set of 15161 Alba domain-containing proteins. Each of these was checked manually for the presence of the Alba domain and compiled into a searchable Shiny R HTML file (**Supplementary Material**).

**Figure 1.**
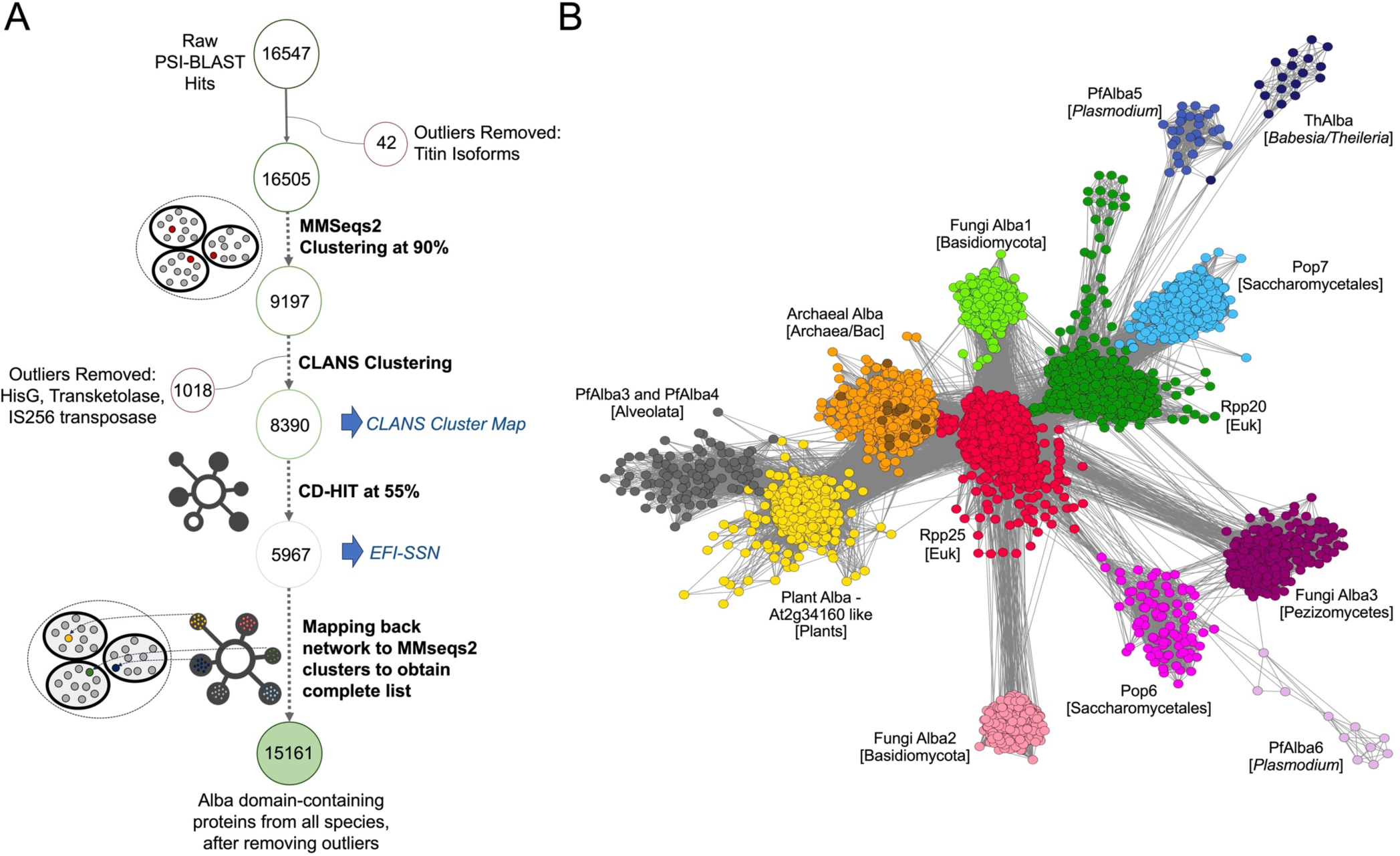
Sequence Similarity Network analysis reveals high diversity within Alba-like proteins. **A.** The analysis workflow of 16547 hits obtained by searching the NCBI protein nr database with 31 Alba seed sequences as queries in PSI-BLAST is shown schematically along with the number of sequences retained after filtering. **B.** A representative sequence similarity network (SSN) of 15161 proteins identified using PSI-BLAST (Altschul et al., 1997) filtered at an E-value threshold of 1E-4. Nodes represent 5967 Alba domain-containing protein sequences clustered using CD-HIT (Huang et al., 2010) at 55% sequence identity. The SSN was generated using EFI-EST (Zallot et al., 2019) and visualized using Cytoscape (Zallot et al., 2019). Each node represents either a single protein or a representative sequence of proteins with similarity greater than 55%. The edge length represents the pairwise sequence identity between two nodes with shorter length indicative of higher sequence identity. Within the Archeal Alba cluster, bacterial nodes are highlighted in brown.

Taxonomic analysis of the 15161 hits indicated that Alba domain-containing proteins are found in all domains of life (**Supplementary Figure S2**), with 55 (0.4%) hits from bacteria, 3862 (25%) hits from archaea, and the remaining 11570 (74%) hits from eukaryotes. An analysis of their amino acid length distribution (**Supplementary Figure S3**) showed that the highest peak containing 1828 sequences was observed for the size range of 90-95 amino acids, which is the average size of the Alba domain in archaea. However, over 56% of the 15161 sequences are greater than 150 amino acids in length, with the longest Alba domain-containing protein containing 4508 amino acids. Given that variations in the amino acid length of homologous proteins from different organisms can represent the diversity of functional roles that they adopt (Lipman et al., 2002), these results suggest that Alba domain-containing proteins may have diverse functions either due to sequence length variations of the Alba domain itself or due to the co-existence of other domains along with the Alba domain. A list of commonly co-occurring domains is provided in **Supplementary Table S1** and includes the Thioredoxin-like_fold, TAXi_N-TAXi_C, HHH_5, TULP and RGG motifs; the latter is found in many RRM domain-containing RNA-binding proteins implicated in neurodegenerative disorders.

### Sequence similarity network analysis reveals the diversity of Alba domain-containing proteins, particularly within unicellular eukaryotes

To obtain a global view of the primary sequence relatedness of the 15161 Alba domain-containing proteins, a sequence similarity network (SSN) was built in the EFI-EST server (Zallot et al., 2019) using a representative set of 5967 sequences, filtered at 55% identity. An alignment score of 4, which corresponded to an e-value threshold of 1E-4, was used to determine the length of 2033164 edges connecting the 5967 nodes. The resulting SSN was visualised in Cytoscape (Shannon et al., 2003) using the prefuse force directed layout with 10000 iterations. As shown in **Figure 1B**, all Alba domain-containing proteins map to 13 unique clusters which correspond to archaeal Albas, plant Alba-like proteins, Rpp20-like or Rpp25-like proteins, PfAlba3/PfAlba4-like proteins, *Plasmodium* Alba5*, Plasmodium* Alba6, Alba-like proteins from *Babesia* and *Theileria* species, *Saccharomycetales* Pop6- or Pop7-like proteins, and three clusters containing fungal Alba-like proteins from Basidiomycota or Ascomycota.

Delving into the features of the SSN, we observed that the largest clusters are formed by the archaeal Albas, Rpp25-like proteins and plant Alba-like proteins (Figure 1B; orange, red and yellow, respectively). The archaeal Alba cluster contains both archaeal and bacterial proteins, with the latter corresponding to proteins either from *Candidatus* uncultured bacteria or from metagenome assemblies (**Supplementary Table S2**). Connected to the archaeal Alba cluster are clusters containing Rpp25, Rpp20 or plant Alba-like proteins (**Figure 1B**) suggestive of the independent evolution of these Alba homologs from the ancient archaeal Alba domain. Interestingly, the Rpp25 cluster (**Figure 1B**; red) contains the largest number of sequences and occupies a pivotal position in the network, forming edges with a majority of the clusters. It contains proteins that are annotated to be Rpp25 in several uni- and multi-cellular eukaryotes, along with PfAlba1 and PfAlba2 (**Supplementary Table S2**). This is in keeping with published reports that have suggested that the Alba domains of PfAlba1 and PfAlba2 are homologous to Rpp25 (Aravind et al., 2003). To our surprise, Pop6, which is thought to be the *S. cerevisiae* Rpp25 homolog, forms an independent cluster (**Figure 1B**; pink) containing other Pop6 proteins from *Saccharomycetales* (**Supplementary Table S2**), with multiple edges connecting it to the Rpp25 cluster. This hints at sequence divergence between Rpp25 and *Saccharomyces* Pop6. Furthermore, three novel clusters, Fungi Alba1, Fungi Alba2 and Fungi Alba3 (**Figure 1B**; fluorescent green, peach and magenta, respectively) formed edges primarily with the Rpp25 cluster and contained uncharacterised or unannotated Alba domain proteins exclusively from fungi. Two of these, Fungi Alba1 and Fungi Alba2, include proteins from Basidiomycota species (**Supplementary Table S2**), while the third, Fungi Alba3, contains proteins from the class Pezizomycetes within Ascomycota (**Supplementary Table S2**). Another cluster that intriguingly connects only with the Pop6 and Fungi Alba3 clusters is constituted by PfAlba6 and its homologs from *Plasmodium* species (**Figure 1B**; mauve) suggestive of a *Plasmodium-*specific divergence/expansion of the Alba domain and that PfAlba6 may perform Pop6-like functions in *Plasmodia*.

The fourth largest cluster is formed by proteins from eumetazoa and fungal species that belong to the Rpp20 family (**Figure 1B**; green) and does not contain any proteins from plants (**Supplementary Table S2**). It forms edges with the archaeal Alba and Rpp25 clusters, in addition to three other clusters, which are described below. In the first cluster (**Figure 1B**; sky blue), we find *Saccharomycetales* proteins (**Supplementary Table S2**) that are similar to *S. cerevisiae* Pop7, which has been described as a homolog of Rpp20 in previous studies (Aravind et al., 2003); this independent Pop7 clustering is similar to the behaviour of the Pop6 cluster relative to Rpp25. The two other Rpp20-connected clusters are the PfAlba5 and ThAlba clusters; the former contains PfAlba5 homologs from *Plasmodium* species (**Figure 1B**; blue; **Supplementary Table S2**) while the latter contains hypothetical proteins from *Babesia* and *Theileria* species (**Figure 1B**; dark blue; **Supplementary Table S2**). This, again, hints at a species-specific evolution of these distant archaeal Alba relatives from Rpp20-like proteins.

Lastly, the plant Alba-like cluster (**Figure 1B**; yellow) is comprised majorly of Alba domain proteins from Viridiplantae (Supplementary Table 1) and a small subset from protists such as Discoban parasites. It forms edges with the archaeal Alba cluster as well as with Rpp25 and Rpp20. In contrast, PfAlba3, PfAlba4, and their homologs from the superphylum Alveolata, cluster together (**Figure 1B**; gray), and only form edges with the plant Alba-like cluster. This was surprising because previous phylogenetic studies had classified PfAlba3 and PfAlba4 as belonging to the Rpp20 family (Aravind et al., 2003; Goyal et al., 2016). Indeed, given the similarity between the PfAlba5 and Rpp20 clusters in our SSN, we hypothesize that, in *P. falciparum,* the functions of Rpp20 might be carried out by PfAlba5 and not by PfAlba3 and PfAlba4.

### Identification of distantly related Alba fold-containing proteins using Hidden Markov Models

Because PSI-BLAST relies on primary sequence similarity for its searches, it cannot identify divergent protein sequences that adopt the Alba fold. Therefore, we decided to use a Hidden Markov Model (HMM)-based approach to identify all proteins that adopt the Alba fold by implementing Hhpred searches in the MPI bioinformatics toolkit (Gabler et al., 2020) against the PDB (Berman et al., 2000), SCOPe (Chandonia et al., 2022; Fox et al., 2014), Pfam (Mistry et al., 2021) and ECOD (Cheng et al., 2014) databases; the 31 seed sequences listed in **Table 1** served as queries. Through this analysis, in addition to known Alba domain-containing proteins, we identified three distant Alba fold-containing proteins: the bacterial sporulation protein VS (SpoVS), the RNA-binding protein YhbY, and the translation initiation factor 3-C (IF3-C). Of these, YhbY and IF3-C have previously been reported to adopt folds similar to the Alba domain (Aravind et al., 2003; Rigden & Galperin, 2008) whereas there is less evidence for SpoVS, although it is annotated as a member of the Alba superfamily in the InterPro database (Paysan-Lafosse et al., 2023). In fact, the Alba fold is thought to have evolved within the archaeo-eukaryotic lineage from the YhbY fold (Aravind et al., 2003).

To verify that SpoVS truly adopts the Alba fold, we quantified the pairwise sequence similarity between the 31 seed Alba sequences and YhbY, IF3-C and SpoVS, using the HH-align module in HH-suite3 of the MPI bioinformatics toolkit. We also quantified the pairwise similarity of their profile HMMs based on Hhpred probability percentages. When these values were visualized using a heatmap (**Figure 2**; left panel), we observed that the highest level of primary sequence similarity for the 31 seed sequences is observed at the SSN cluster level, for example, between the archaeal and bacterial Alba proteins, amongst the Alba domain-containing proteins from multicellular eukaryotes, and so on. This was expected since SSN clustering is based on primary sequence identity. In contrast, the distantly related Alba fold-containing proteins, including SpoVS, showed 0.1 to 7.9% sequence similarity to the 31 seed Alba domain proteins and this is probably why they were not identified by PSI-BLAST. Next, upon comparison of the Hhpred probability percentages (**Figure 2**; right panel), amongst the Alba fold-containing proteins, SpoVS showed a high Hhpred probability percentage (greater than 90%) with most of the seed Alba sequences, which indicates that it adopts a fold very similar to the Alba domain and can be confidently classified as a member of the Alba superfamily. Contrarily, bacterial YhbY showed a maximum Hhpred probability percentage of 23% with an archaeal Alba, *Aeropyrum pernix* Alba 1, and *Escherichia coli* IF3-C did not show significant HMM profile similarity with any of the proteins, including YhbY and SpoVS.

**Figure 2.**
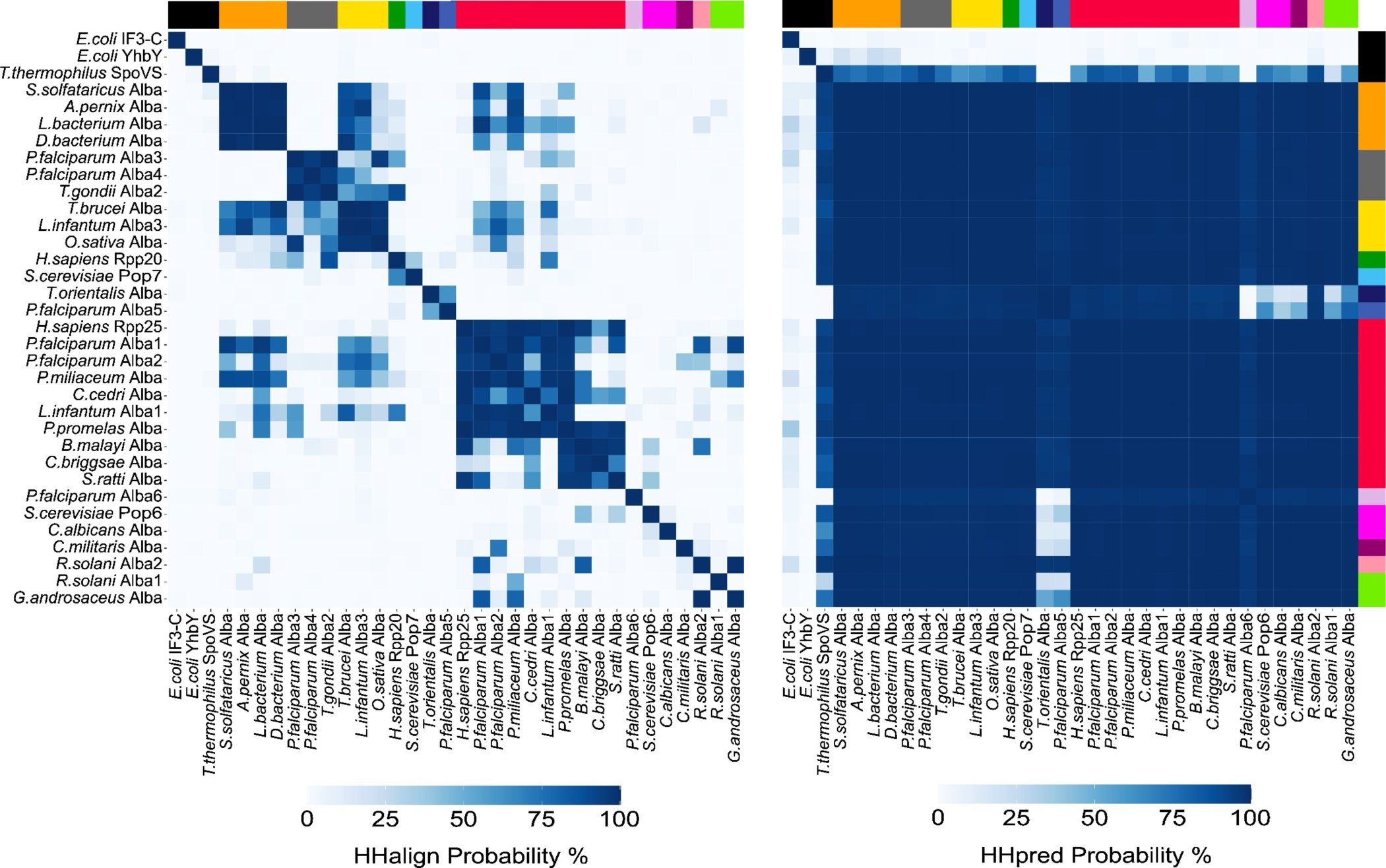
HMM profiles reveal the similarity between Alba domain-containing proteins and SpoVS, a member of the Alba superfamily. Pairwise comparison of primary amino acid sequence (***left panel***) and HMM profiles (***right panel***) of the Alba domain of 31 seed Albas (refer to Table 1 for more details) and three Alba fold-containing proteins, *T. thermophilus* SpoVS (PDB ID:2EK0), *E. coli* YhbY (PDB ID:1LN4) and *E. coli* IF3-C (PDB ID:2IFE). The colour bar represents the SSN cluster to which a given sequence maps to, with black representing the distant homologs identified using Hhpred. Hhalign probability and Hhpred probability values were calculated using the Hhalign module in HH-suite3 (Steinegger et al., 2019).

Focusing on the *P. falciparum* Alba proteins, the Alba domains of PfAlba1 and PfAlba2 shared sequence similarity percentages greater than 50% with Rpp25 and most other Alba domain-containing proteins, be it archaeal, bacterial or eukaryotic, but not with PfAlba3, PfAlba4, PfAlba5, PfAlba6, or select fungal Alba-like proteins that belong to the Fungi Alba2 and Fungi Alba3 clusters (**Figure 2**; left panel). PfAlba3 showed a sequence similarity percentage greater than 50% only with PfAlba4, with Alba domains from select unicellular parasitic eukaryotes, and with the Alba-like protein of *Oryza sativa*, while PfAlba4 showed a sequence similarity percentage greater than 50% only with PfAlba3 and with Alba domain-containing proteins of *Trypanosoma brucei brucei*. Interestingly, PfAlba5 shared a sequence similarity percentage greater than 50% only with *Theileria* and *Babesia* Albas while PfAlba6 did not show sequence similarity percentages greater than 50% with any of the proteins included in this analysis. The pattern changed dramatically when we looked at Hhpred probability percentages (**Figure 2**; right panel): for PfAlba1, PfAlba2, PfAlba3 and PfAlba4, HMM profile similarity with the remaining Alba seed sequences as well as with SpoVS increased to greater than 90%. For PfAlba5, the probability percentages with most of the seed Albas were high, except for PfAlba6, Pop6, representative sequences from the Fungi Alba1 and Fungi Alba3 clusters, and SpoVS. PfAlba6 showed the most striking shift: although sequence similarities with the other seed Albas were low, HMM profile similarities were high. In fact, the lowest probability percentages were with PfAlba5, *Theileria* Alba and SpoVS. Overall, we infer that although there is low primary sequence conservation amongst Alba domain proteins, they largely adopt the same three-dimensional fold, which can be further extrapolated to all of the 15161 Alba hits identified in this study.

### Structural analysis of Alba domain-containing proteins reveals deviations within the newly identified SSN clusters

Further, to explore tertiary structure similarity, we used RoseTTAFold (Baek et al., 2021), an ‘accurate’ structure prediction algorithm, to predict the Alba domain structures of the 31 seed Alba sequences and the three distant Alba fold-containing proteins; where available, we also included crystal structures in this analysis. As shown in **Figure 3A**, RoseTTAFold was able to predict the Alba domain structures of all seed sequences with high confidence, except for the *Brugya malayi* Alba protein which belongs to the Rpp25 cluster; hence, this structure was not included for further analysis. To assess structural prediction accuracy, the RoseTTAFold-predicted structures of archaeal *Sulfolobus solfotaricus* Alba, *Homo sapiens* Rpp25, *H. sapiens* Rpp20, *S. cerevisiae* Pop6 and *S. cerevisiae* Pop7 were compared to their respective crystal structures using TM-align (C. Zhang et al., 2022). The resulting TM-align scores were greater than 0.9 indicating high levels of tertiary structure identity (**Supplementary Figure S4**).

**Figure 3:**
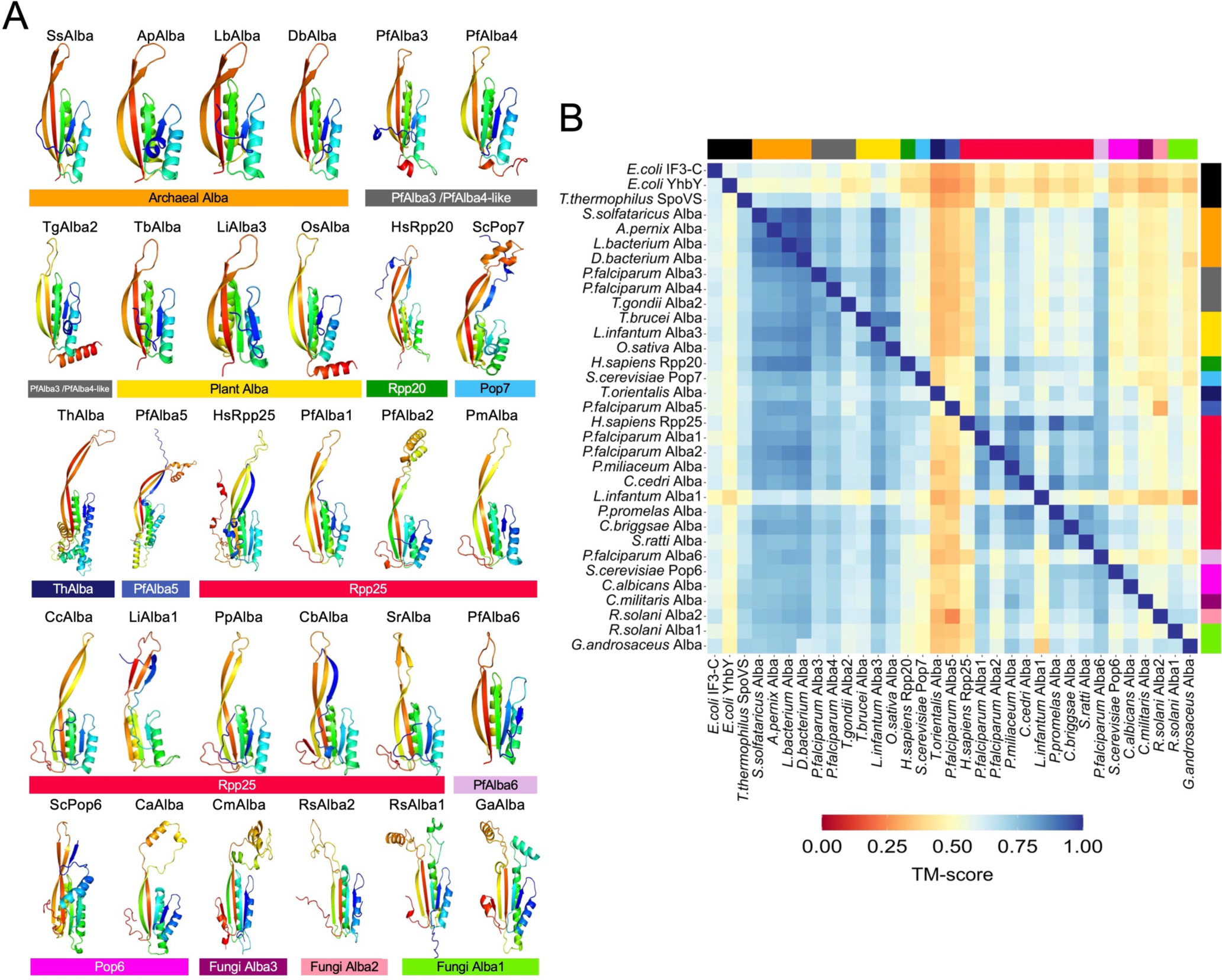
Structural analysis of the Alba-like proteins. **A.** Structures of the Alba domain of representative SEED sequences from each cluster were predicted using RoseTTAFold (Baek, Minkyung, et al. 2021 (12)). The color bar represents the cluster to which each structure belongs. **B.** Quantitative pairwise comparisons of the 31 representative Alba domain structures and structures of distant homologs were performed using TM-align (Zhang et al., 2005). TM-scores of 0.0–0.30 indicate random structural similarity; TM-scores of 0.5–1.00 indicate that the two proteins adopt generally the same fold.

We thereafter scrutinized the remaining Alba domain structures. At a first glance, all predicted seed Alba structures revealed conservation of the classic Alba fold: a short N-terminal β-sheet followed by an α-helix, a second β-sheet, another α-helix, and two elongated β-sheets at the C-terminus joined by a flexible loop, *i.e,* β1−α1−β2−α2−β3−β4 (**Figure 3A**). A few interesting deviations are highlighted here: (i) When it comes to lower eukaryotic species, Alba domains have elongated C-terminal β-sheets. A specific case is PfAlba5 which is predicted to contain β3 and β4 of length 21-22 amino acids as compared to archaeal Albas which contain 11-15 amino acid-long β3 and β4. (ii) *S. cerevisiae* Pop6 and Pop7 diverge from Rpp25 and Rpp20, respectively, through the presence of an extra α-helix and β-sheet at the N-terminus (**Figure 3A**). (iii) The Alba domains of PfAlba3, PfAlba4 and plant Albas contain an extra C-terminal α-helix which is absent in Rpp20, further validating that PfAlba3 and PfAlba4 are not homologs of Rpp20 and may have unique functions in *P. falciparum*. (iv) The newly identified fungal Alba proteins (**Figure 1B**; fluorescent green, peach and magenta clusters) appear to have extra β-sheets and α-helices between β3−β4 which loop out of the Alba fold. However, because these are predicted structures, we cannot conclusively comment on this particular structural variation and its implications.

Subsequently, in a pairwise manner, we compared the RoseTTAFold-predicted structures of the seed Albas to each other or to the crystal structures of SpoVS, YhbY and IF3-C using TM-align (**Figure 3B**). The TM-align scores of all Alba domain structures relative to the archaeal Alba domain were greater than 0.7, indicating strong tertiary structure identity (**Figure 3B**). However, the pariwise TM-align scores for some of the other comparisons were lesser than 0.7, for instance when PfAlba5 and *Theileria* Alba were used as the reference sequence for comparison. Notably, for all seed Albas, the TM-align scores relative to SpoVS, YhbY and IF3-C ranged from 0.4 to 0.6, indicating the common topology that these proteins adopt, despite having very little sequence similarity. This supports the evolutionary relatedness of SpoVS, YhbY and IF3-C to the Alba domain. Lastly, the examination of the electrostatic potential surface map of the predicted Alba seed structures revealed the presence of a highly electropositive pocket, across the board, that most likely corresponds to the site of nucleic acid binding (**Supplementary Figure S5**).

### The Alba dimer interface shows the highest amino acid sequence conservation

We next looked for sequence conservation amongst all 15161 Alba domain-containing proteins identified in this study relative to archaeal Alba using Scorecons (Valdar, 2002). Additionally, to infer the functional relevance of conservation, we mapped these amino acid residues onto the crystal structure of archaeal Alba(s) in the presence or absence of DNA and/or RNA (PDB IDs:2BKY, 6LT7 and 3IAB (Jelinska et al., 2005; Perederina et al., 2007; Yin et al., 2021)). As seen in **Figures 4A and 4B**, for the 15161 Alba domain proteins, β2, α2 and β4 contain a higher number of conserved residues, a majority of which are important for dimerization and to a lesser extent, nucleic acid interaction. This pattern of conservation is also observed within the archaeal Alba cluster (**Figure 4C**). Of note, the Gly43 residue of archaeal Alba, known for its involvement in RNA binding (L. Guo et al., 2014), showed significant conservation in all Albas. In contrast, the Lys16 residue, subject to acetylation in archaeal Alba and a purported regulator of DNA binding activity (Bell et al., 2002), exhibited poor conservation (**Figures 4A and 4C**).

**Figure 4.**
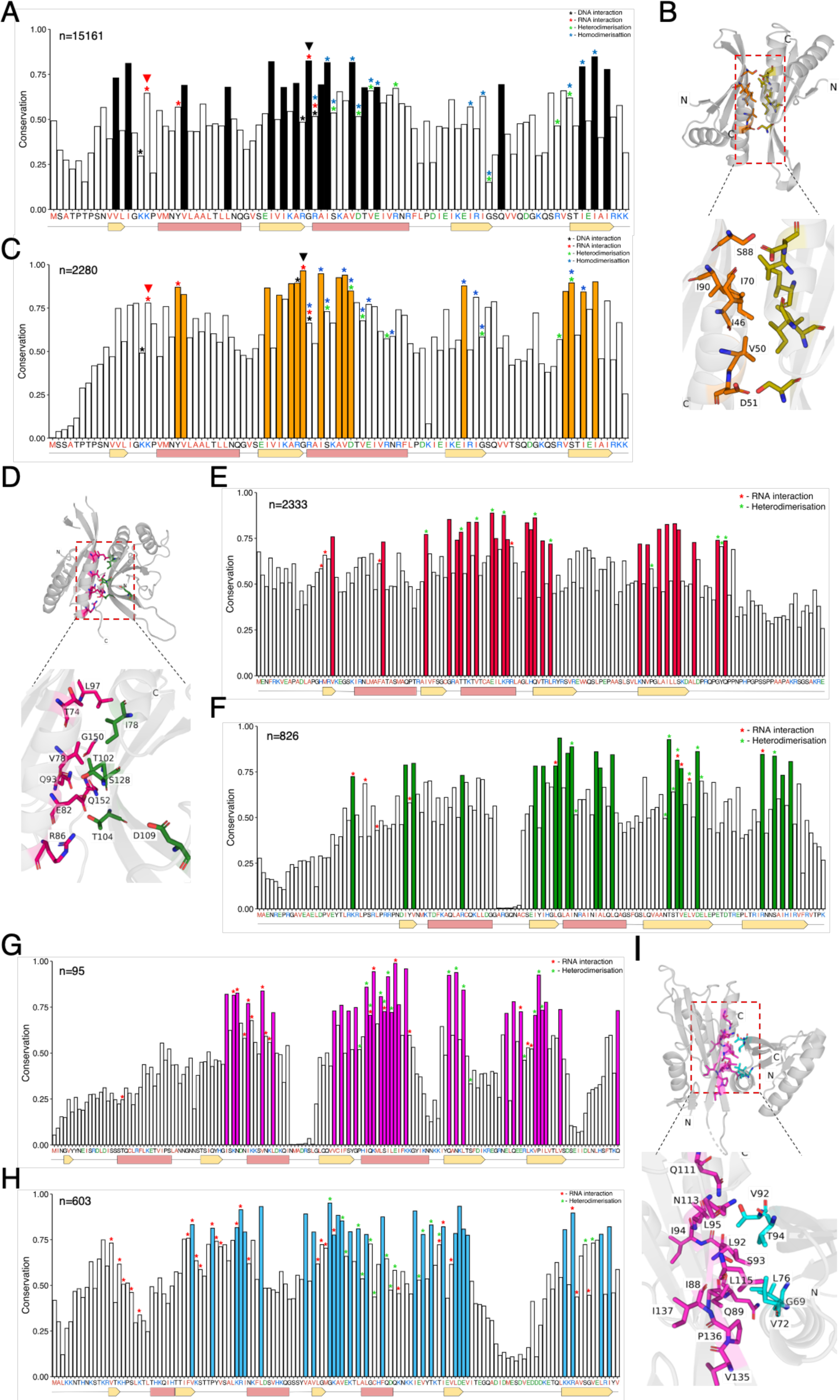
Residues at the dimer interface of Alba homo-/hetero-dimers are significantly conserved. **A**. Amino acid sequence conservation of all 15161 Alba homologs was measured relative to an archaeal Alba sequence (Alba1 of *S. solfotaricus*), represented along the X-axis. **B**. Crystal structure of the dimer interface of the Alba1:Alba2 heterodimer from *S. solfotaricus* (PDB: 2BKY (Jelinska et al., 2005)) is shown, with key residues highlighted. **C.** Conservation of 2280 sequences present in the archaeal Alba cluster was measured relative to Alba1 of *S. solfotaricus*, represented along the X-axis. **D.** Crystal structure of the dimer interface of human Rpp20-Rpp25 in complex with the P3 stem loop of RNase MRP RNA (PDB: 6LT7 (Yin et al., 2021)) is shown. **E & F.** Amino acid conservation of protein sequences present in the **(E)** Rpp25 (n=2333) and **(F)** Rpp20 (n=826) clusters, respectively. The human homologs are represented along the X-axis. **G & H.** Amino acid conservation of the proteins present in the **(G)** Pop6 (n=95) and **(H)** Pop7 (n=603) clusters, respectively. The *S. cerevisiae* homologs are represented along the X-axis. **I.** Crystal structure of the dimer interface of *S. cerevisiae* Pop6/Pop7 in complex with the P3 region of the RNase MRP RNA (PDB: 3IAB (Perederina et al., 2007)) is shown. For parts A, B, E, F, G and H, conserved residues of at least one standard deviation more than the mean conservation across the alignment are shown as colour-filled bars. *highlights functionality of select residues as per the key provided in the figure. In the consensus sequence (X-axis), red represents hydrophobic residues, blue represents positively charged residues, and green represents negatively charged residues. The protein secondary structure is plotted along the X-axis with α−helices coloured in pink and β−sheets in yellow as per the corresponding crystal structure.

Given the availability of RNA-bound crystal structures of the Rpp20-Rpp25 and Pop6/Pop7 heterodimers, we further looked at sequence conservation within these sub-groups and mapped conserved residues onto the crystal structure. In the Rpp20-Rpp25 heterodimer (PDB: 6LT7 (Yin et al., 2021)), we observed a similar pattern as above, wherein the residues responsible for heterodimerization exhibited a high degree of conservation along with the equivalent of archaeal Alba Gly43 (**Figures 4D-F**). Conversely, in the case of the Pop6/Pop7 complex (PDB: 3IAB (Perederina et al., 2007)), residues involved in both RNA interaction and heterodimerization display significant conservation (**Figures 4G-I**).

Lastly, sequence conservation analysis for Alba clusters lacking crystal structures (**Supplementary Figure S6**) revealed that in the PfAlba3/PfAlba4 cluster, only β2, including the glycine corresponding to archaeal Gly43, exhibits high conservation, while within the plant Alba cluster, β2 and α2 show high conservation, but not β4. Remarkably, conservation scoring of the Fungi Alba2 cluster resulted in the detection of a putative Tubby-like domain (TULP) in conjunction with the Alba domain in all 380 members (**Supplementary Figure S6**). The TULP domain is postulated to act as a transcription factor (Carroll et al., 2004), and may diversify the functionality of its associated Alba domain. Taken together, our analysis suggests that the dimer interface – for either homo- or hetero-dimerization – is the most highly conserved for experimentally characterized Albas. This can be extrapolated to understand the functionality of the newly identified Alba domain-containing proteins and the impact of homo- or hetero-dimerization on protein function.

### Phylogenetic analysis of Alba-like proteins

Thus far, SSN analysis has provided a comprehensive understanding of the inter-relatedness amongst Alba homologs at the primary sequence level, while secondary and tertiary structure analyses have revealed structure-function relationships for the 13 Alba sub-groups. To obtain insights into evolutionary relatedness, we constructed a maximum-likelihood phylogenetic tree using a selected set of 4666 representative sequences from the 13 SSN-derived Alba clusters, including all 31 seeds. Because Alba sequences are extremely divergent, with pairwise sequence identity as little as 10%, we were unable to find a suitable outgroup and hence, we built an unrooted tree (**Figure 5**). The branching of the tree independently supported the clustering observed in SSN analysis, with each major Alba subgroup demonstrating a significant branching probability. Nonetheless, a few deviations were evident, which are discussed below.

Considering the archaeal Albas to be the most ancient members of this protein family, we observed a single branchpoint from which the Rpp20- and Rpp25-like clades diverged (**Figure 5**; marked with an arrowhead). Within the former, the earliest diverging branch was comprised of PfAlba5-like proteins and the ThAlbas. Given that these proteins formed separate clusters in the SSN (**Figure 1B**; blue and dark blue), we were surprised to find that they were not distinguishable phylogenetically. The second diverging branch in the Rpp20-like clade was comprised of PfAlba3/PfAlba4-like proteins and plant Albas, while the third branch included Rpp20 proteins from fungi, metazoa as well as fungal homologs of Pop7, as separate groups. To a large extent, these branches reflected the clustering observed in the SSN; the only exception was the divergence of fungal and metazoan Rpp20 homologs. On the other hand, within the Rpp25-like clade, the earliest diverging branch was comprised of PfAlba1/PfAlba2 homologs from the Stramenopiles-Alveolata-Rhizaria or SAR taxonomic group followed by distinct branches containing plant and metazoan Rpp25-like proteins (**Figure 5**), all of which constituted the red cluster in the SSN of **Figure 1B**. This was unexpected and we attributed it to the identity threshold of 55% that we had chosen for SSN clustering. Indeed, when the SSN threshold was increased to 65% identity, we observed that PfAlba1/PfAlba2 and the plant Rpp25-like proteins formed independent clusters which shared edges with a metazoan Rpp25 cluster (data not shown). Other notable features of the phylogenetic tree include the diversification of Pop6-like and PfAlba6-like proteins from a common ancestor within the Rpp25-like clade, branching of the newly identified fungal Albas from a single node in the Rpp25-like clade, and the lack of a single branch for bacterial Albas within the archaeal clade. This latter observation supports the acquisition of Albas by bacteria through horizontal gene transfer. Overall, phylogenetic analysis provided a timeline for the genetic divergence of the Alba domain and hints at potential time periods when new functions were gained.

**Figure 5:**
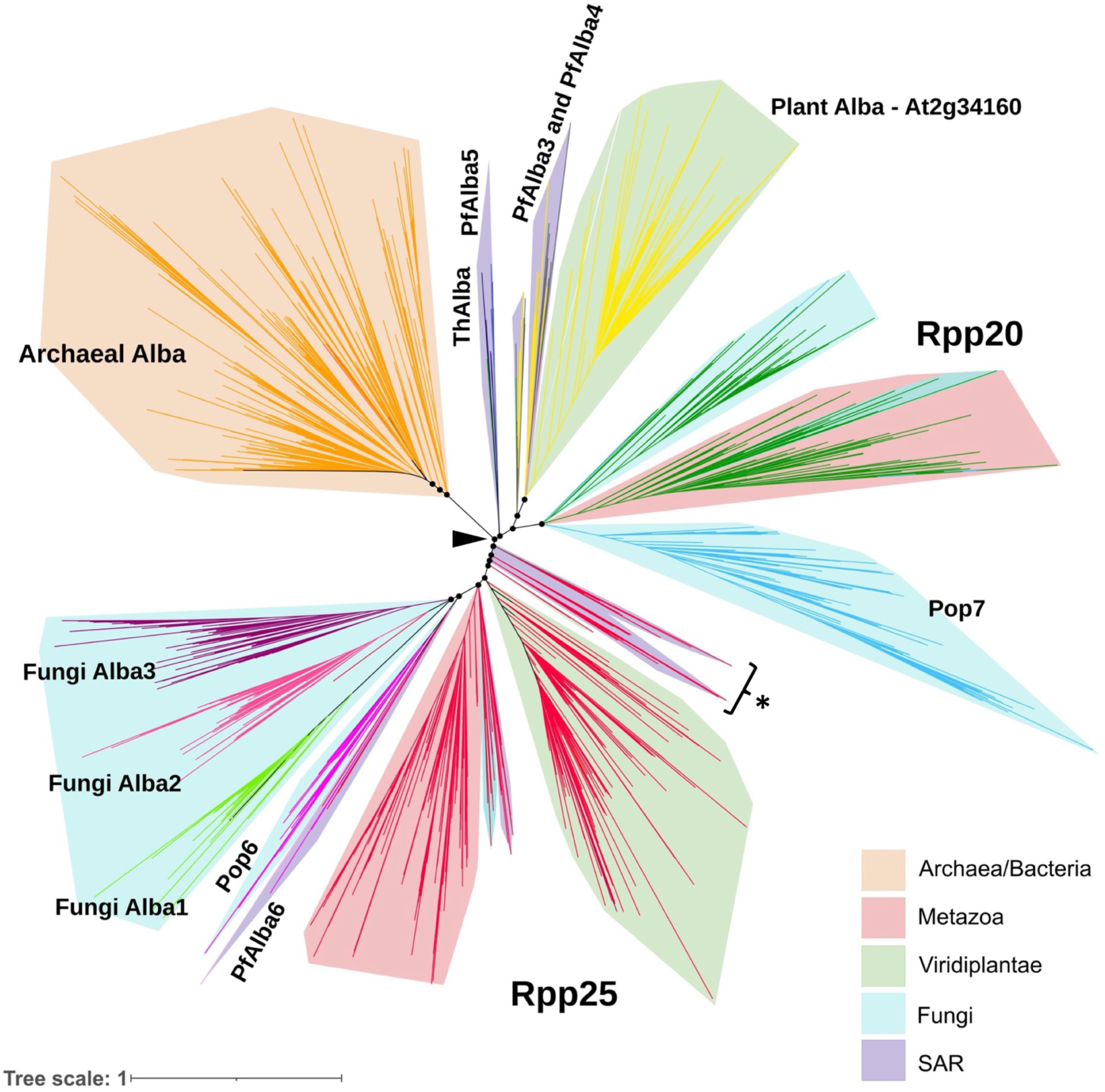
Phylogenetic reconstruction of Alba relatedness. A maximum likelihood tree was built using a representative set of 4666 Alba homologs, which were obtained by clustering the 15161 hits at a pairwise sequence identity threshold of 70%. Only branch points with an aBayes branch support value greater than 0.75 are represented, resulting in the retention of 4585 Alba homologs in the final tree. The branches are colored based on the cluster information from SSN analysis. Coloured ranges depict different kingdoms. The arrowhead denotes the branchpoint for divergence of Rpp20- and Rpp25-like proteins. }* highlights the branch containing PfAlba1 and PfAlba2. SAR = Supergroup of Protists containing Stramenopiles, Alveolata and Rhizaria.

## Discussion

For a given protein domain, amino acid sequences change over time to adapt to physiological and environmental stresses. These changes could either be single amino acid substitutions, additions and deletions, or the incorporation of small peptide units into the basic protein scaffold, all of which increase functionality and provide a fitness benefit (Alva et al., 2015; Anantharaman et al., 2001; Ponting & Russell, 2002). Nonetheless, despite the accumulation of changes, it has been observed that a given domain largely retains its three-dimensional functional fold, even in evolutionarily divergent organisms. To understand this phenomenon in detail, researchers have begun performing in depth studies of sequence, structural and functional relationships of ubiquitous proteins, domains and/or folds such as the histone fold, SH3 and OB domains in ribosomal proteins, P-loop NTPases and Rossmann-fold enzymes, glycosyltransferases, ESCRT systems, etc. (Alva & Lupas, 2019; Alvarez-Carreño et al., 2021; Grau-Bové et al., 2022; Longo et al., 2020; Makarova et al., 2024; Taujale et al., 2020), in turn tracing their evolutionary origin. But, analyses of more specialized protein domains lag behind. Here, we use a combination of exhaustive database searches, SSN creation, structural comparison, and phylogenetic reconstruction to understand the inter-relatedness and evolutionary dynamics of the DNA/RNA-binding Alba domain, highlighting its versatility and importance to essential biological processes across all kingdoms of life.

Firstly, by building an SSN containing more than 15000 Alba domain-containing proteins from different taxonomic lineages, we identified 13 distinct Alba clusters, with eight eukaryotic clusters specific to taxonomic groups such as the genus *Plasmodium*, order Saccharomycetales or division Basidiomycota. Subsequent phylogenetic analysis provided an evolutionary framework for these groupings: for instance, in the SSN, the archaeal Albas clustered with a very small set of Alba domain proteins from bacteria, amongst which 13% mapped to the phylum Proteobacteria and 27% to the Terrabacteria clade, which includes the ancient phyla Actinobacteria and Firmicutes. Moreover, because a majority of the bacterial hits were derived from metagenome assemblies – *i.e.,* their taxonomic classification contains a level of uncertainty – we postulated that select bacteria may have acquired Alba proteins from archaea through horizontal gene transfer. This was supported by the phylogenetic tree, wherein the bacterial Albas did not fall into a single branch but were spread across different branches within the archaeal clade. On the whole, we can irrefutably conclude that this DNA/RNA-binding domain was not present in the Last Universal Common Ancestor, thus upholding a seminal study which arrived at a similar conclusion by comparing fewer than 70 archaeal Alba, eukaryotic Rpp20-like, and eukaryotic Rpp25-like protein sequences (Aravind et al., 2003). Of note, the Alba domain of Macronuclear development protein 2 or Mdp2 from the ciliate *Stylonychia lemnae*, which was used by Aravind *et al*. to benchmark Rpp25-like Alba proteins but was not included in our list of Alba seed sequences, falls within the Rpp25-like cluster of the SSN and phylogenetic tree.

Another striking outcome of SSN and phylogenetic analyses was the identification of lineage-specific diversification of eukaryotic Albas. In fact, 9 of the 13 SSN clusters corresponded to protist and fungal taxonomic groups, suggesting that these unicellular organisms may have diversified their Alba repertoire to regulate DNA- and RNA-dependent processes required for specific lifestyle adaptations such as parasitism. A case in point is the malaria parasite which contains six Albas mapping to four SSN clusters: Rpp25-like (containing PfAlba1 and PfAlba2), PfAlba3/PfAlba4-like, *Plasmodium* Alba5 and *Plasmodium* Alba6, the latter two clusters being genus-specific. Phylogenetic analysis further segregated PfAlba1 and PfAlba2 into the SAR clade, *i.e.,* based on taxonomic grouping within Protista, and away from plant and metazoan Rpp25 proteins, suggesting that PfAlba1 and PfAlba2 functions may have diverged away from RNase P/MRP regulation. This is evidenced by their correlation to mRNA translation regulation and *Plasmodium* virulence gene expression (Acharya et al., 2024; Chêne et al., 2012; Diffendall et al., 2023; Goyal et al., 2012; Vembar et al., 2015). Nonetheless, the question remains, what is the nature of the RNase P/MRP complex in *Plasmodium*? In humans and budding yeast, it is well-established that Rpp25/Pop6 and Rpp20/Pop7 carry out similar functions in the context of the RNase P/MRP complex during 5’ tRNA processing (Chamberlain et al., 1998; Frank & Pace, 1998; Goldfarb & Cech, 2017; Guerrier-Takada et al., 2002; Jarrous, 2017; Welting et al., 2006). However, in *A. thaliana* and *T. brucei*, the ribonucleoprotein RNase P complex is substituted by a proteinaceous RNase P, PRORP, for tRNA processing (Bhatta & Hillen, 2022). Whether *Plasmodium* employs a similar strategy or whether PfAlba5 and PfAlba6 take on the roles of Rpp20/Pop7 and Rpp25/Pop6, respectively, remains to be seen. Another interesting case is a parasite closely related to *Plasmodium, T. gondii.* The four Alba-domain containing proteins of this parasite segregate into four distinct groups: Rpp25-like (TgAlba1), Pop7 (ToxoDB ID/NCBI ID: TGME49_245520/XP_018634887), Rpp20-like (ToxoDB ID/NCBI ID: TGME49_217665/XP_018634862), and PfAlba3/PfAlba4-like (TgAlba2). Experimental characterization of TgAlba1 and TgAlba2 showed that both proteins co-localize *in vivo* and are involved in translational control of gene expression (Gissot et al., 2013). Whether they function as a heterodimer is not known, neither is the function of the remaining TgAlbas.

Secondly, sequence conservation analysis across the 13 SSN clusters did not identify a single invariant amino acid residue that participated in nucleic acid recognition. Instead, dimerization emerged as a universal feature of the Alba protein family, substantiated by the high conservation of the dimer interface. This has been experimentally demonstrated. In archaea, the Alba1:Alba2 heterodimer showed a greater level of DNA compaction as compared to the homodimer of either Alba (Jelinska et al., 2005) (Laursen et al., 2021). Thus, by differentially regulating the expression of Alba1 and Alba2, chromatin compaction could be modulated, in turn altering gene expression. In humans, the Rpp20-Rpp25 heterodimer functions as a single unit, with heterodimerization being a prerequisite for interaction with the P3 RNA of RNAse P (Hands-Taylor et al., 2010). In yeast, a similar prerequisite for Pop6-Pop7 heterodimer formation has been reported (Houser-Scott et al., 2002; Perederina et al., 2007) In *A. thaliana*, AtAlba1 and AtAlba2, which cluster with the plant Alba-like family in the SSN, bind to genic R-loops as heterodimers and maintain genome stability (Yuan et al., 2019). Lastly, in addition to heterodimerization, homodimerization has also been demonstrated for select Albas *in vitro* Chêne et al., 2012; Nag et al., 2024; Wardleworth et al., 2002). Taken together, these studies suggest that Alba proteins recognise bipartite motifs in nucleic acids. Moreover, in organisms that express more than two Alba proteins, the permutations of heterodimers that can form may increase dramatically, which could impart divergent functions to this protein family.

At the global level, all of the 15161 proteins that we obtained through our exhaustive searches adopted the β1−α1−β2−α2−β3−β4 Alba fold. However, the lack of any invariant residues across their Alba domains indicates the extensive primary sequence innovations that this domain has undergone. A protein fold is said to be ‘innnovable’ when the structural scaffold, which provides robustness and stability to the fold, and the active site residues, which recognise substrates, are highly separated (Tóth-Petróczy & Tawfik, 2014); this enables substrate diversification, which can consequently impart diverse functions. In the case of the Alba domain, the scaffold remains highly conserved, thus retaining a highly electropositive surface for nucleic acid binding. However, by varying the sequence as well as length of loops between the structural elements, the domain may have gained the ability to recognise different types of nucleic acids and sequence motifs. We further speculate that other molecules with electronegative surfaces such as RNA-binding proteins with intrinsically disordered regions could also be recognised. Indeed, the innovability of this domain can be observed in organisms such as *Plasmodium* where there is lineage-specific evolution, and also in the new fungal Albas, which have a characteristic extension of α2 and new loops in between β3 and β4.

Collectively, our findings strongly indicate the evolutionary trajectory of Alba proteins, transitioning from sequence-independent NAPs in archaea, primarily responsible for genomic organization, to specialized proteins that exhibit selective recognition of specific nucleic acids in a particular protein complex or pathway. Notably, nucleic acid binding activity has been retained across all members of the Alba superfamily, underscoring its pivotal role in diverse biological contexts. This can be correlated to other protein families such as bacterial integration host factor or IHF, a homolog of histone HU which evolved to become a sequence-specific transcriptional activator at certain genetic loci while maintaining genome organization capability at others (Browning et al., 2010). Another example is the chromatin architectural protein TK0471 from the archaea *Thermococci kodakarensis,* which was identified to be homologous to the TrmB transcription factor, a specialised regulator of the *mal* operon (Maruyama et al., 2011). Lastly, two archaeal proteins, Smj12 and Abfr1, which non-specifically bind to and condense DNA (Li et al., 2017; Napoli et al., 2001), are both members of the Lrs14 family of specialised transcription factors (Orell et al., 2013).

Overall, our sequence-structure analysis serves as a starting point for inferring the functionality of a range of Alba proteins. If we consider that proteins which perform the same function most likely share similar sequences, and hence cluster together, then we can assign specific functions to select Alba clusters. Additionally, we can predict the heterodimer pairs that could form in organisms with three or more Alba domain proteins. For example, the primary functions of proteins in the Rpp25-like, Rpp20-like, Pop6 and Pop7 clusters is most likely RNase P/MRP activity, with additional moonlighting functions such as the regulation of telomerase activity; the resulting heterodimer pairs could be arrived at in a similar fashion. On the other hand, the major functions of proteins in the plant Alba-like cluster, which contains entries from plants as well as Discoba, could be translational regulation and maintenance of genome stability. Extrapolating further, all proteins in the PfAlba3/PfAlba4-like cluster may possess site-driven endonuclease activity in the presence of divalent metal ions *in vitro*, as was recently demonstrated for PfAlba3 (Banerjee et al., 2023), although the *in vivo* significance of nuclease activity remains to be explored. Nonetheless, we cannot rule out that the function of a given Alba domain-containing protein may be determined not just by the Alba domain, but by its co-occurring domains, post-translational modifications, and/or sub-cellular localisation. Further studies of individual Albas from diverse clusters and clades will be required to dissect out the range of cellular processes controlled by this protein family.

In conclusion, our investigation into the evolutionary dynamics and functional diversity of the Alba protein domain sheds light on its remarkable adaptability and significance across the biological spectrum, particularly in the context of genome maintenance and RNA biology. Moreover, our study provides a foundational framework for predicting functional roles and potential heterodimerization partners within the Alba protein family, paving the way for future investigations into the cellular processes controlled by this protein domain. To the best of our knowledge, this study is the first of its kind for a specialised DNA/RNA/DNA:RNA-binding domain, and paves the way for similar studies into innovable protein domains.

## Acknowledgements

This work was supported by the Ramalingaswami Re-entry Fellowship Contingency Grant (No. BT/RLF/Re-entry36/2017) awarded by the Department of Biotechnology, India, to S.S.V. Work in the S.S.V. lab is supported by a grant-in-aid to IBAB from the Department of IT, BT, S&T of the Government of Karnataka, India. We acknowledge Vikram Alva of the Max Planck Institute for Developmental Biology, Tubingen, Germany, for scientific discussions, and Shubhada Hegde of IBAB, Bengaluru, India, for critical reading of the manuscript.

## Author contributions

J.J. and S.S.V. conceived and designed the experiments; J.J. performed all of the experiments; J.J. and S.S.V. analyzed the data and performed statistical analyses; J.J. and S.S.V. wrote the paper; All authors read and approved the final manuscript.

