## Supplementary Material for "Sequence, Structural and Functional Diversity of the Ubiquitous DNA/RNA-Binding Alba Domain"

**Supplementary Information for Jaiganesh and Vembar, 2024**

*This document contains:*

**Supplementary Tables S1 and S2**

**Supplementary Figures S1 to S6**

*Provided as a link:*

**Shiny R app summarising the Alba-domain proteins identified in this study with their NCBI accession ID and the SSN cluster that they belong to**

**http://plasmobees.shinyapps.io/Alba_domain_containing_proteins**

**Supplementary Table S1:** Summary of co-occurring domains in Alba domain-containing proteins. Domains that co-occur in a minimum of five Alba hits are listed below.

| Domain Name(s) | Domain Function(s) | InterPro/PFAM ID(s) |
|---|---|---|
| Thioredoxin-like_fold | Regulates the activity of target proteins by changing the redox state of thiol groups (S2 to SH2) | IPR012336 |
| TAXi_N; TAXi_C | Aspartic proteinase and xylanase inhibitory activity | IPR032861; IPR032799 |
| HHH_5 | Helix-hairpin-helix DNA-binding domain | PF14520 |
| TULP | Tubby-like domain thought to have transcription factor activity | IPR000007 |
| PCI_dom; 26S_Psome_reg_C | Proteasome, COP9, Initiation factor 3 or PCI domain stabilizes protein-protein interactions and co-occurs with the 26S_Psome_reg_C domain, which is found at the C-terminus of 26S proteasome regulatory subunits. | IPR000717; IPR013586 |
| DCTN3 | Dynactin subunit 3, which may play a role in cytokinesis | IPR009991 |
| LMBR1-like_membr_prot | LMBR1-like membrane protein, which may play a role in cell signaling | IPR006876 |
| PFD_beta-like | Prefoldin beta-like; protein chaperone | IPR002777 |
| PsdUridine_synth_N | Pseudouridine synthase II, N-terminal; catalyse the isomerisation of uridine to pseudouridine (Psi) in a variety of RNA molecules. | IPR002501 |
| KH_dom_type_1 | K Homology domain, type 1 is a RNA-binding domain found in a variety of proteins that regulate eseential processes such as splicing, etc. | IPR004088 |
| Cyt_c_oxidase_su5A/6 | Subunits Va and VI are the heme A-containing chains of cytochrome c oxidase. Subunit Va is located on the matrix side of the membrane and binds thyroid hormone T2, releasing allosteric inhibition caused by the binding of ATP to subunit IV and allowing high turnover at elevated intramitochondrial ATP/ADP ratios. | IPR003204 |
| 5'(3')-deoxyribonucleotidase | Not known | IPR010708 |

**Supplementary Table S2:** Summary of the 13 SSN clusters identified in this study and the number of Alba domain-containing proteins belonging to each cluster. The interactive Krona charts are deposited in https://github.com/jaiganeshjg/Alba as html files and can be accessed by clicking the provided hyperlink.

| Cluster | Number of sequences | Krona chart |
|---|---|---|
| Alba domain-containing proteins | 15161 | https://raw.githack.com/jaiganeshjg/Alba/main/All_Alba.tsv.krona.html |
| Rpp25 | 5095 | https://raw.githack.com/jaiganeshjg/Alba/main/Rpp25.tsv.krona.html |
| Archaeal Alba | 3913 | https://raw.githack.com/jaiganeshjg/Alba/main/ArchaealAlba.tsv.krona.html |
| Plant Alba | 1781 | https://raw.githack.com/jaiganeshjg/Alba/main/PlantAlba%E2%80%93At2g34160like.tsv.krona.html |
| Rpp20 | 1460 | https://raw.githack.com/jaiganeshjg/Alba/main/Rpp20.tsv.krona.html |
| Pop7 | 1034 | https://raw.githack.com/jaiganeshjg/Alba/main/Pop7.tsv.krona.html |
| Fungi alba3 | 676 | https://raw.githack.com/jaiganeshjg/Alba/main/FungiAlba3.tsv.krona.html |
| Fungi alba2 | 433 | https://raw.githack.com/jaiganeshjg/Alba/main/FungiAlba2.tsv.krona.html |
| Fungi alba1 | 360 | https://raw.githack.com/jaiganeshjg/Alba/main/FungiAlba1.tsv.krona.html |
| PfAlba3 and PfAlba4 | 152 | https://raw.githack.com/jaiganeshjg/Alba/main/PfAlba3_and_PfAlba4.tsv.krona.html |
| Pop6 | 145 | https://raw.githack.com/jaiganeshjg/Alba/main/Pop6.tsv.krona.html |
| PfAlba5 | 53 | https://raw.githack.com/jaiganeshjg/Alba/main/PfAlba5.tsv.krona.html |
| PfAlba6 | 39 | https://raw.githack.com/jaiganeshjg/Alba/main/PfAlba6.tsv.krona.html |
| ThAlba | 20 | https://raw.githack.com/jaiganeshjg/Alba/main/ThAlba.tsv.krona.html |

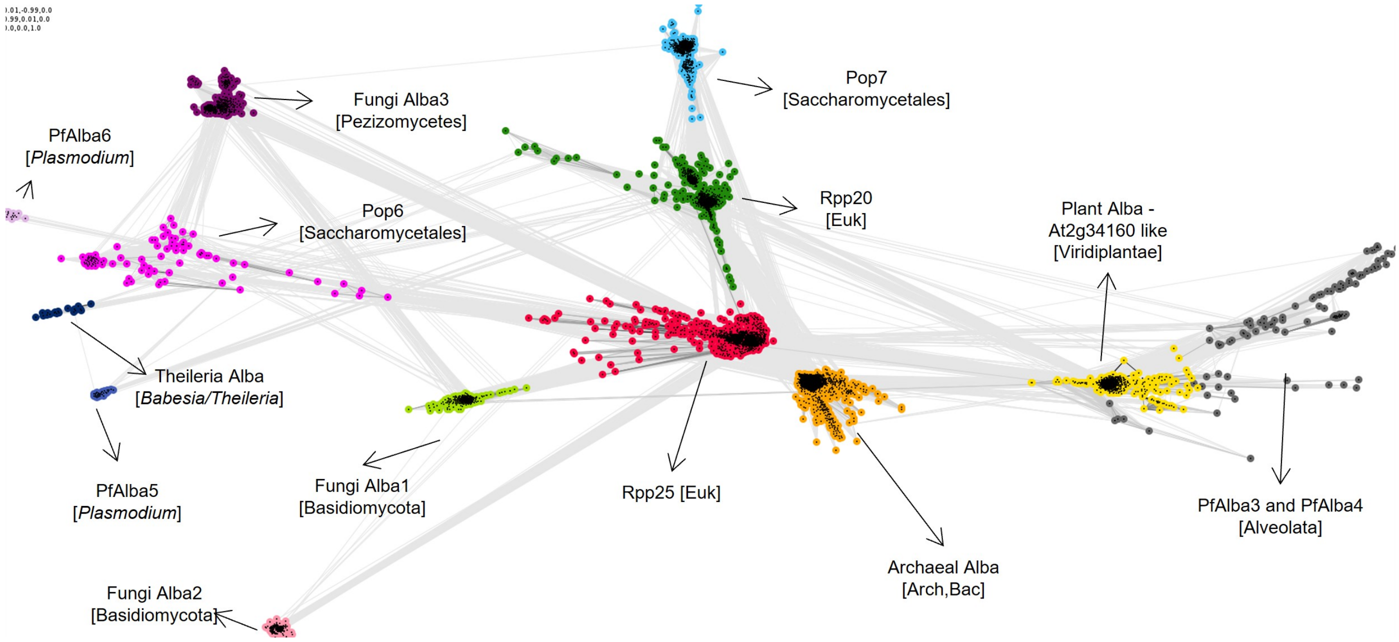


**Supplementary Figure S1. CLANS Cluster map of a representative set of 8390 Alba homologs** obtained after clustering the 15161 Albas identified in this study at a maximum pairwise identity of 90% using MMseqs2 (Steinegger et al., 2021). Dots represent sequences which were clustered at a p-value cutoff of 1e-4. Sequences belonging to the same cluster are similarly coloured. The degree of their pairwise sequence similarities, as measured by BLAST+ p-values, is reflected in line colouring; the darker a line, the greater the similarity.

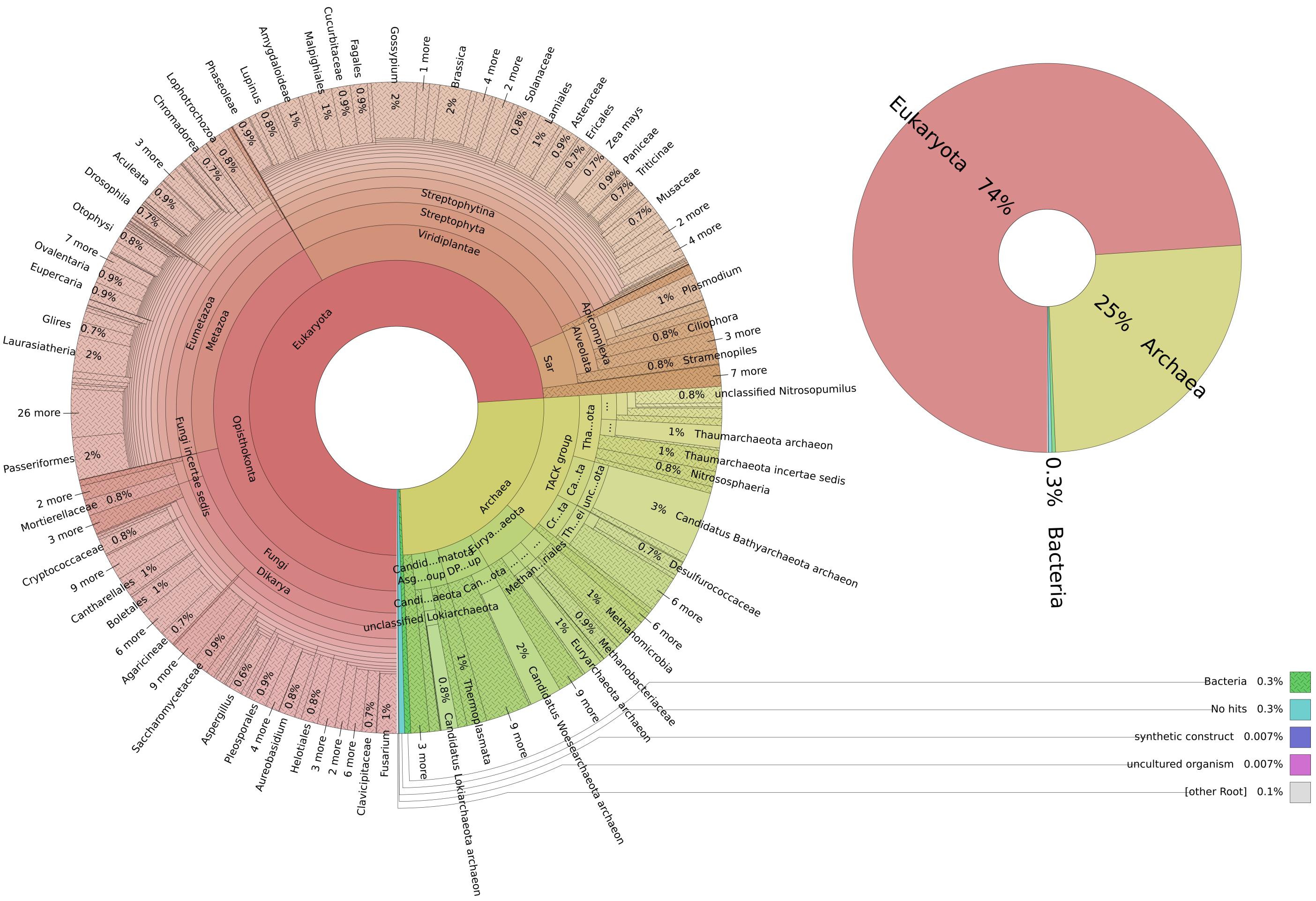


**Supplementary Figure S2. Taxonomic distribution of 15161 Alba domain-containing proteins.** ***Left***: A pie chart was generated in Krona (Ondov et al., 2011) to represent the taxonomic distribution of all 15161 Alba domain-containing proteins identified in this study. The innermost rings represent domain and kingdom level classification. Proceeding outwards, the taxonomic resolution increases, *i.e.,* Phylum, Class, Order, Family, Genus, and Species being the outermost ring. ***Right***: A Krona pie-chart showing domain-level taxonomic distribution of the 15161 Alba homologs. The Krona charts are available as interactive html format. Refer to Supplementary Table S1 for more details.

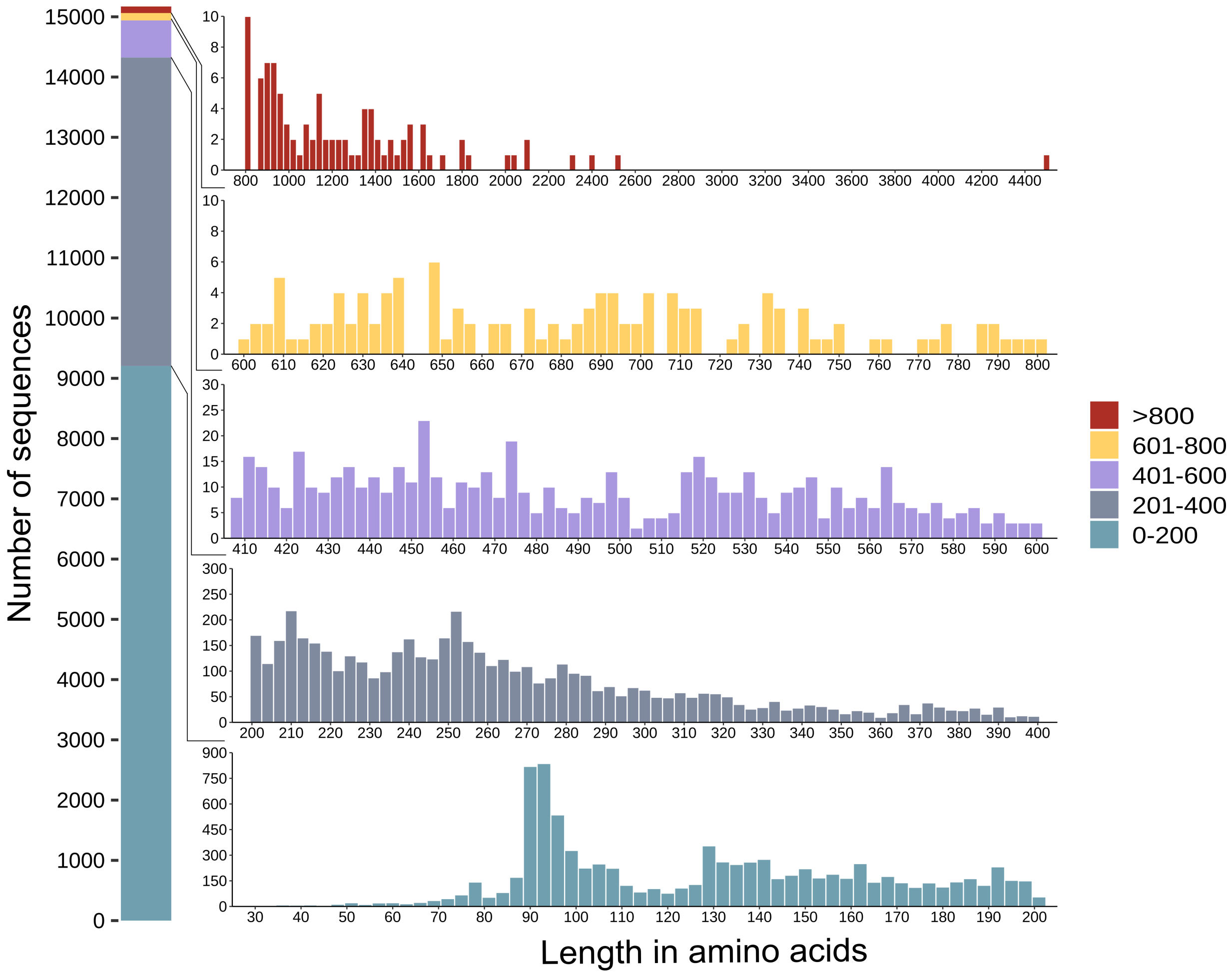


**Supplementary Figure S3. Amino acid sequence length distribution of 15161 Alba-domain containing proteins.** The X-axis corresponds to the length of the protein in amino acids, and the Y-axis to the number of hits for a given length.

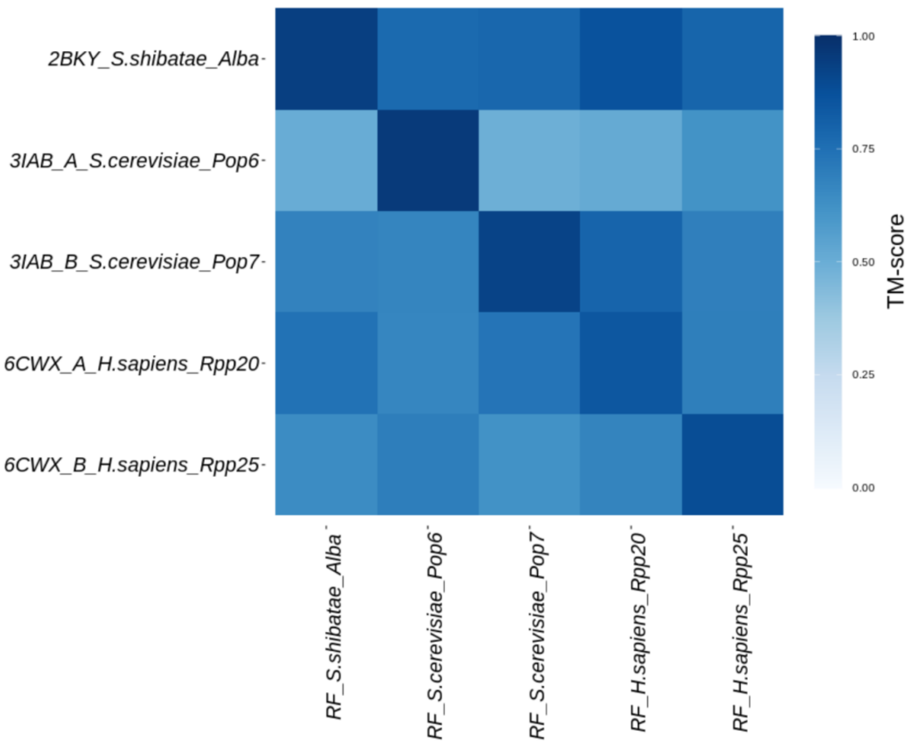


**Supplementary Figure S4. Quantitative pairwise comparison of the solved crystal structures and RoseTTAFold-predicted structures of select Alba domains**. TM-align (Zhang et al., 2022) was used to compare the crystal structures (y-axis) of five Albas to their *ab initio* structures (x-axis) predicted using RoseTTAFold (Baek et al., 2021). Crystal structures were downloaded from the RCSB-PDB database and their PDB ID is provided. RF = RosettaFold. The scale of the heatmap shows the TM-score reported by TM-align.

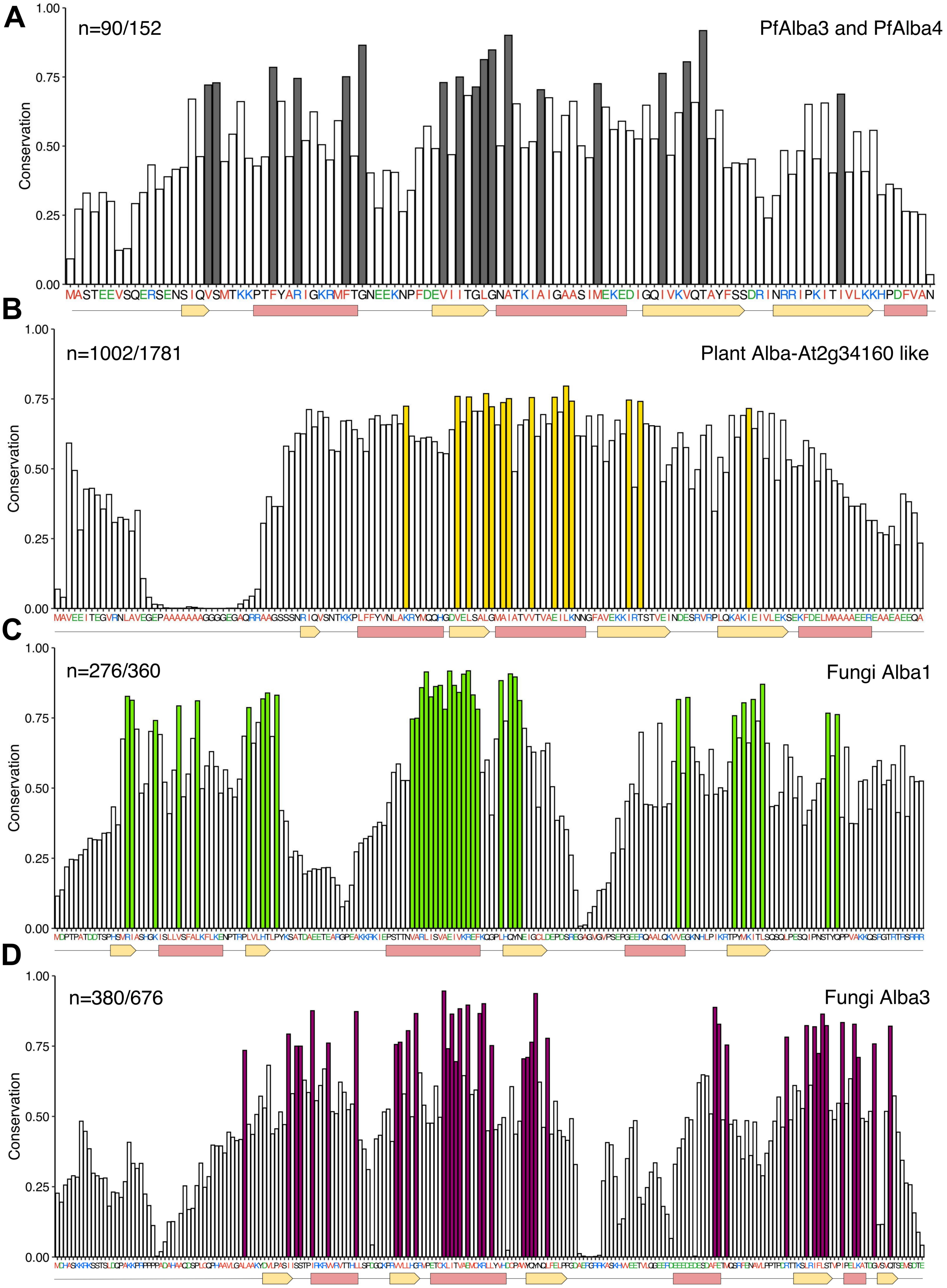
A
n=90/152
PfAlba3 and PfAlba4
Conservation
1.00
0.75
0.50
0.25
0.00
MASTEEVSQERSENSIQVSMTKKPTFYARIGKRMFTGNEEKNPFDEVIITGLGNATKIAIGAASIMEKEDIGQIVKVQTAYFSSDRINRRIPKITIVLKKHPDFVAN
B
n=1002/1781
Plant Alba-At2g34160 like
Conservation
C
n=276/360
Fungi Alba1
Conservation
D
n=380/676
Fungi Alba3
Conservation

E

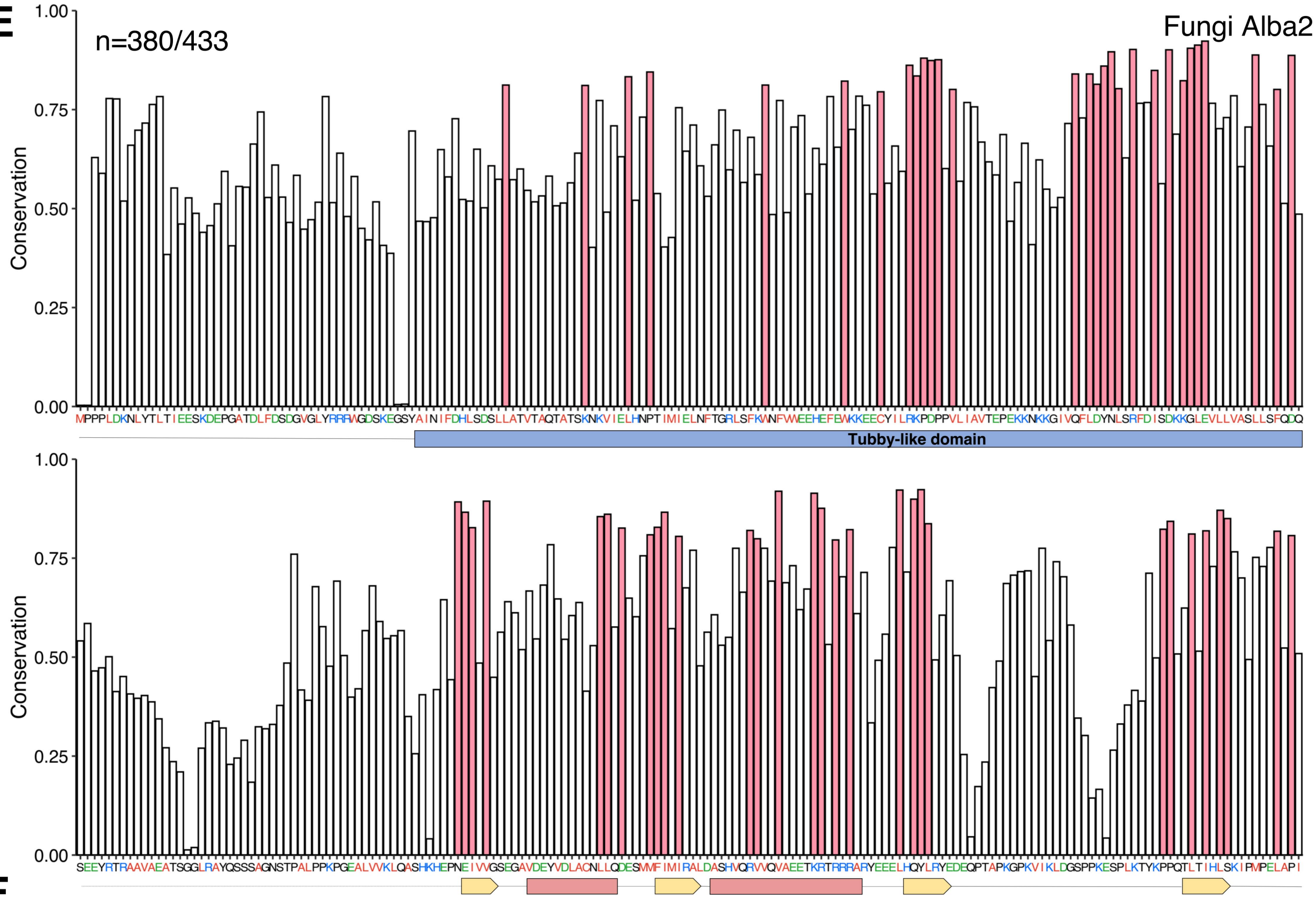
n=380/433
Fungi Alba2
Conservation
1.00
0.75
0.50
0.25
0.00
Tubby-like domain

F

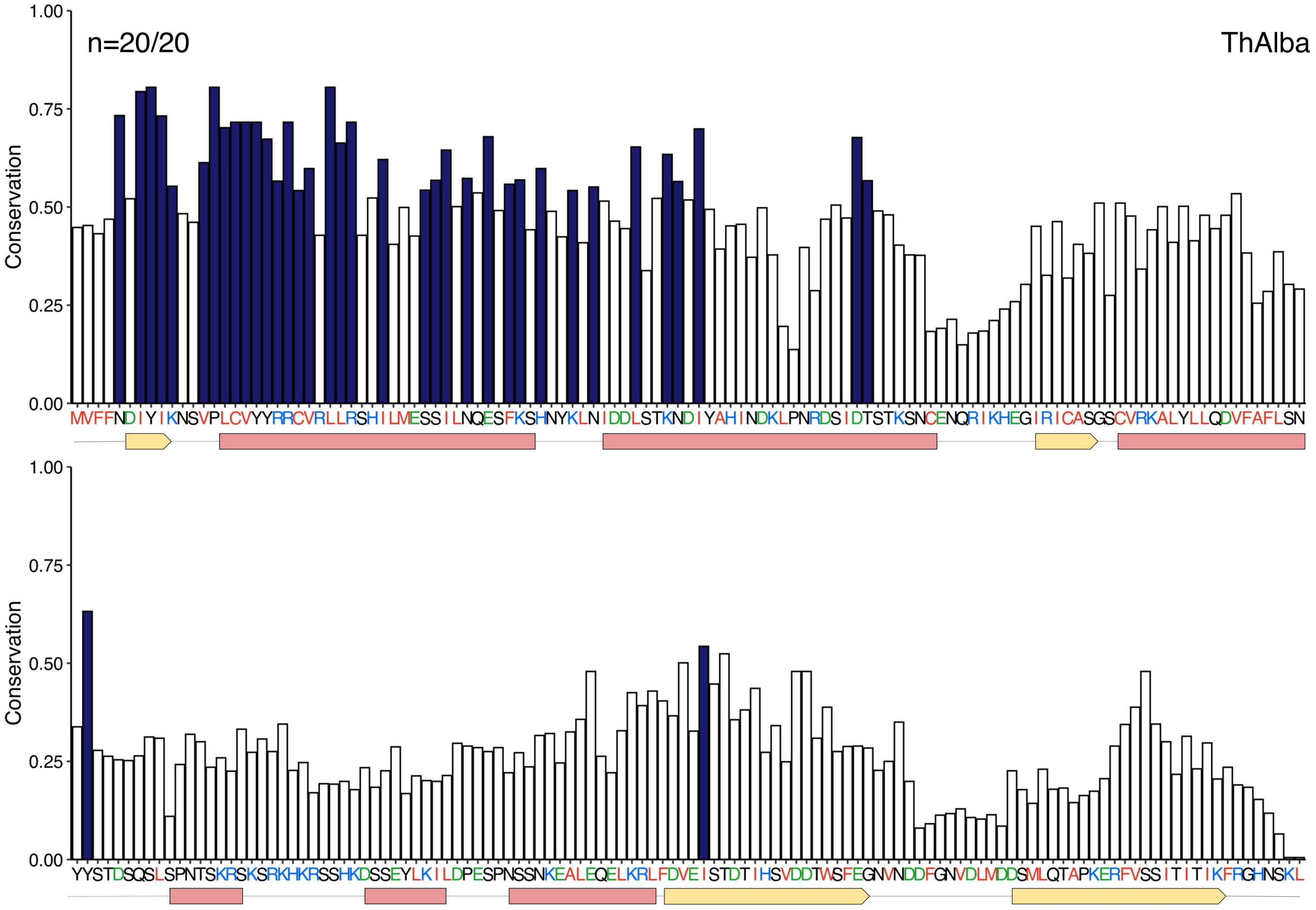
n=20/20
ThAlba
Conservation
1.00
0.75
0.50
0.25
0.00

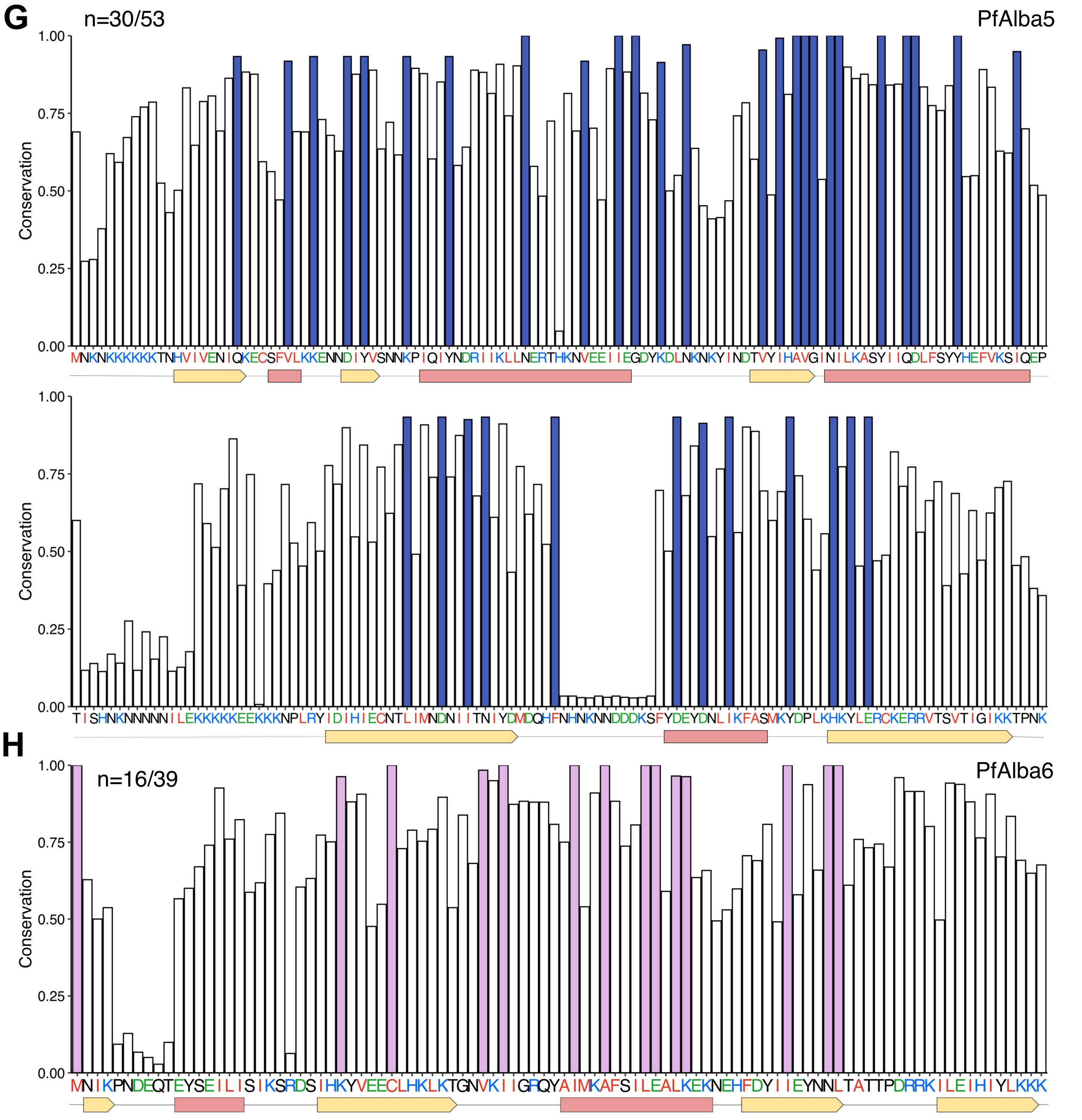


**Supplementary Figure 5. Sequence Conservation of Alba proteins from eight different SSN clusters.** The conservation (Y-axis) of the indicated number of proteins from, **A.** PfAlba3/PfAlba4-like cluster; **B.** Plant Alba-like cluster**;** **C.** Fungi Alba1 cluster; **D.** Fungi Alba3 cluster; **E.** Fungi Alba2 cluster; **F.** Alba-like proteins from *Babesia* and *Thileria* species; **G.** *Plasmodium* Alba5 cluster; or **H.** *Plasmodium* Alba6 cluster, was scored against the 'seed' sequence of that particular cluster using the ScoreCons server (Valdar, 2002). The number of aligned sequences out of the total number of sequences is shown. Conserved residues of at least one standard deviation more than the mean conservation across the alignment are shown as colour-filled bars, and their colour corresponds to the cluster colour in Fig. 1B. In the consensus sequence (X-axis), red represents hydrophobic residues, blue represents positively charged residues, and green represents negatively charged residues. The protein secondary structure is plotted along the X-axis with $\alpha$-helices coloured in pink and $\beta$-sheets in yellow.

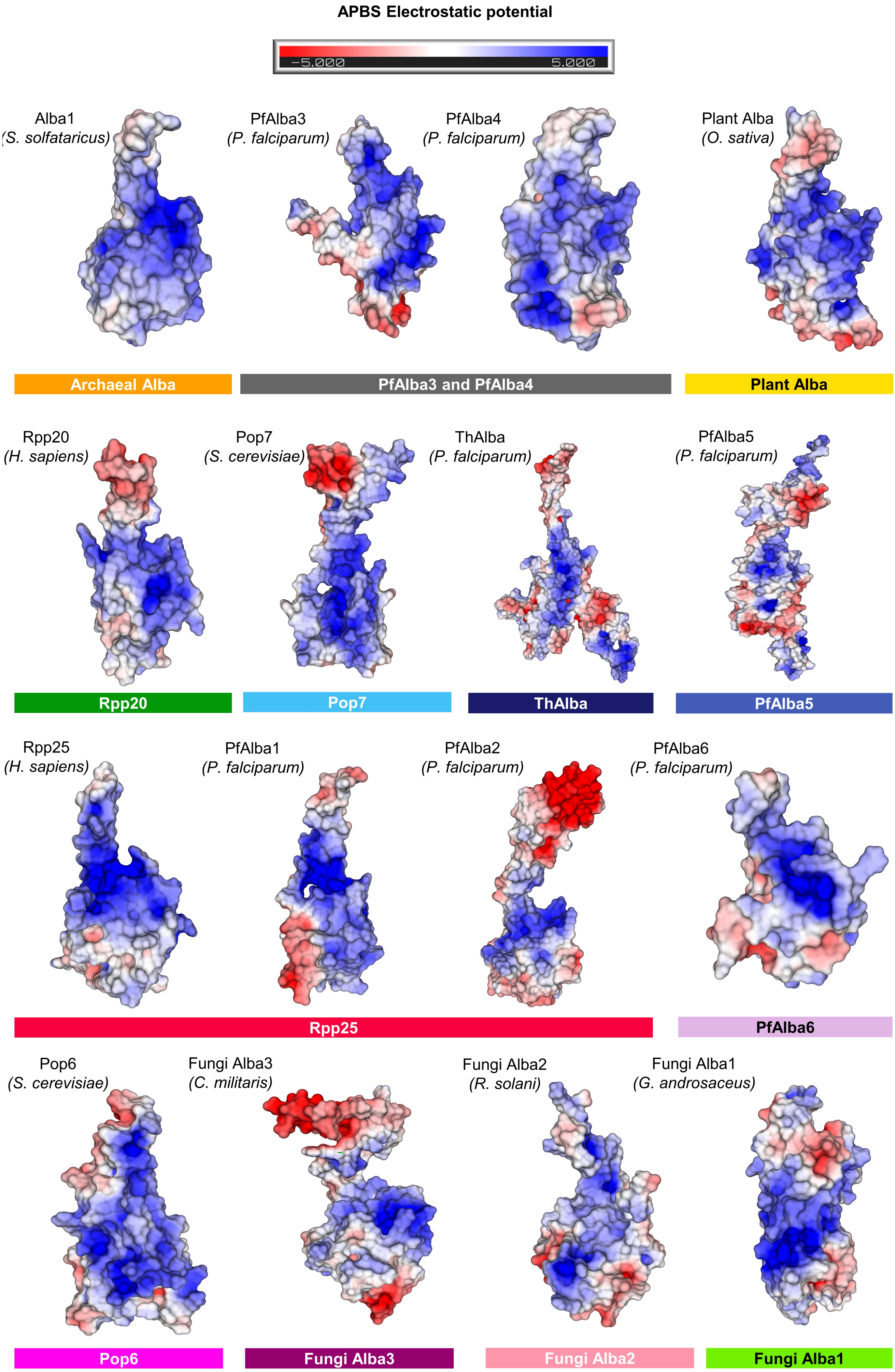
APBS Electrostatic potential
-5.000
5.000
Alba1
(S. solfataricus)
PfAlba3
(P. falciparum)
PfAlba4
(P. falciparum)
Plant Alba
(O. sativa)
Archaeal Alba
PfAlba3 and PfAlba4
Plant Alba
Rpp20
(H. sapiens)
Pop7
(S. cerevisiae)
ThAlba
(P. falciparum)
PfAlba5
(P. falciparum)
Rpp20
Pop7
ThAlba
PfAlba5
Rpp25
(H. sapiens)
PfAlba1
(P. falciparum)
PfAlba2
(P. falciparum)
PfAlba6
(P. falciparum)
Rpp25
PfAlba6
Pop6
(S. cerevisiae)
Fungi Alba3
(C. militaris)
Fungi Alba2
(R. solani)
Fungi Alba1
(G. androsaceus)
Pop6
Fungi Alba3
Fungi Alba2
Fungi Alba1

**Supplementary Figure 6. The DNA/RNA-interacting electropositive surface is conserved across all Alba homologs.** For one representative structure from each cluster, as well as for the predicted structures of all six *Plasmodium* Albas, Adaptive Poisson-Boltzmann Solver (APBS; (Baker et al., 2001)) surface maps were generated and rendered at +/- 5 kT/e. The scale bar represents charge, with blue indicating electropositivity and red indicating electronegativity.

**List of References**


1. Baek, M., DiMaio, F., Anishchenko, I., Dauparas, J., Ovchinnikov, S., Lee, G. R., Wang, J., Cong, Q., Kinch, L. N., Dustin Schaeffer, R., Millán, C., Park, H., Adams, C., Glassman, C. R., DeGiovanni, A., Pereira, J. H., Rodrigues, A. V., Van Dijk, A. A., Ebrecht, A. C., … Baker, D. (2021). Accurate prediction of protein structures and interactions using a three-track neural network. *Science*, *373*(6557), 871–876.
2. Baker, N. A., Sept, D., Joseph, S., Holst, M. J., & McCammon, J. A. (2001). Electrostatics of nanosystems: Application to microtubules and the ribosome. *Proceedings of the National Academy of Sciences*, *98*(18), 10037–10041. https://doi.org/10.1073/pnas.181342398
3. Ondov, B. D., Bergman, N. H., & Phillippy, A. M. (2011). Interactive metagenomic visualization in a Web browser. *BMC Bioinformatics*, *12*. https://doi.org/10.1186/1471-2105-12-385
4. Steinegger, M., Mirdita, M., Karin, E. L., Von Den Driesch, L., Galiez, C., & Söding, J. (2021). *MMseqs2 User Guide*.
5. Valdar, W. S. J. (2002). Scoring residue conservation. *Proteins*, *48*(2), 227–241. https://doi.org/10.1002/prot.10146
6. Zhang, C., Shine, M., Pyle, A. M., & Zhang, Y. (2022). US-align: Universal structure alignments of proteins, nucleic acids, and macromolecular complexes. *Nature Methods*, *19*(9), 1109–1115. https://doi.org/10.1038/s41592-022-01585-1
